# Druggability of Phospholipase C-β Isoforms with Small Peptides Patterned after the Autoinhibitory XY Linker

**DOI:** 10.1101/2025.07.22.666078

**Authors:** Jorge de Andrés-López, David Cabañero, Isabel Devesa, Shihab Shah, Gregorio Fernández-Ballester, Nikita Gamper, Mª Ángeles Bonache, Rosario González-Muñiz, Asia Fernández-Carvajal, Antonio Ferrer Montiel

**Affiliations:** Instituto de Investigación, Desarrollo e Innovación en Biotecnología Sanitaria de Elche (IDiBE). Universitas Miguel Hernández, Elche, Spain; Faculty of Biological Sciences, School of Biomedical Sciences. University of Leeds, Leeds, United Kingdom; Department of Pharmacology; The Key Laboratory of Neural and Vascular Biology, Ministry of Education; The Key Laboratory of New Drug Pharmacology and Toxicology, Hebei Medical University, Shijiazhuang, Hebei, China; Instituto de Química Médica. Consejo Superior de Investigaciones Científicas (CSIC). Madrid, Spain

**Author notes:** To whom correspondence should be addressed. IDiBE-UMH, Av de la Universidad s/n, 03202 Elche. Spain.

**Keywords:** PLCβ, XY linker, peptide inhibitors, TRPV1, inflammation, nociception, pain, pharmacological tool

## Abstract

The phospholipase C-β (PLCβ) signaling pathway plays a pivotal role in peripheral nociception, particularly during inflammation and pain transduction. Despite their validation as important therapeutic targets, PLCβ isoforms are yet undruggable due to the difficulties to identify potent and selective modulators. Here, we addressed this question and used the autoinhibitory XY linker present in these enzymes as a source of peptide inhibitors of PLCβ activity. We report that peptides patterned after this motif inhibited PIP_2_ hydrolysis and the consequent calcium release from endoplasmic reticulum. In primary nociceptor cultures, active peptides notably attenuated bradykinin-induced electrogenesis and TRPV1 sensitization, thus reducing nociceptor hyperexcitability. Noteworthy, intraplantar administration of a lead peptide prevented inflammation and hypersensitivity in a mouse model of inflammatory pain. Collectively, our findings indicate that peptides patterned after the autoinhibitory XY linker act as selective PLCβ inhibitors with *in vivo* anti-inflammatory and antinociceptive activity, providing pharmacological tools for this enzyme family.

**SIGNIFICANCE:** Phospholipases C (PLC) are intracellular signaling proteins, with PLCβ isoforms crucial in somatosensory neuron signaling. These enzymes interact with G-protein coupled receptors for pro-inflammatory and algesic agents, sensitizing nociceptors by increasing their excitability. Despite their importance, selective PLCβ modulators remain limited; U73122 is widely used, though it lacks specificity and has off-target effects. Here, we introduce peptide inhibitors based on the XY autoinhibitory motif that selectively block PLCβ activity, reduce bradykinin-induced neuronal responses and TRPV1 sensitization, and do not affect other PLC isoforms. In a murine inflammatory pain model, local administration of our lead peptide showed both anti-inflammatory and antinociceptive effects, highlighting its therapeutic potential. This approach expands the toolkit of PLC-isoform selective modulators for drug development.

## INTRODUCTION

Phospholipases C (PLC) catalyze the hydrolysis of phospholipid phosphatidylinositol 4,5-bisphosphate (PIP_2_) into inositol 1,4,5-trisphosphate (IP_3_) and diacylglycerol (DAG). IP_3_ binding to its receptors promotes calcium release from intracellular stores, while DAG activates protein kinases C (PKC). These second messengers trigger a wide range of intracellular events, ranging from ion channel modulation to exocytosis or cell proliferation and differentiation^1–4^. The family of mammalian PLC includes six subtypes of isozymes with different isoforms: β (1-4), γ (1,2), δ (1,3,4), ε, ζ and η (1,2), each one with tissue-specific expression patterns^1^.

PLCβ isoforms are relevant in somatosensory perception of exogenous and endogenous stimuli, and in neurotransmission to the central nervous system. Several studies have proved PLCβ expression, especially the PLCβ3 isoform, in different neuronal subpopulations of dorsal root ganglia (DRG) and trigeminal ganglia^5,6^. In these cells, PLCβ isoforms modulate neuronal excitability through G-protein coupled receptors (GPCRs), as they are differentially activated by Gα_q_ and Gβγ subunits^7,8^. Interestingly, the PLCβ pathway modulates Transient Receptor Potential (TRP) channel activity^9^. This type of Ca^2+^-permeable non-selective cation channels work as molecular sensors of thermal, mechanical and chemical stimuli^10^. By far, the best-characterized member is the vanilloid receptor 1 (TRPV1), gated by the vanilloid compound capsaicin, noxious heat (> 43°C), acidic pH or depolarizing voltages^11^. TRPV1 is expressed in subpopulations of Aδ and C-fibers, where it is a key detector and transducer of painful stimuli^12,13^.

Upon injury, peripheral tissues release pro-inflammatory mediators and sensitize nociceptors, leading to hypersensitivity and inflammatory pain^14^. These agents activate multiple sensitizing mechanisms, which are orchestrated by GPCRs-PLCβ pathways. A prominent example is TRPV1 sensitization^15,16^. Activation of PKCε downstream of PLCβ phosphorylates serine residues in TRPV1, lowering the activation threshold to its activating stimuli^17,18^. This effect synergizes with the PLCβ-induced reduction in PIP_2_ levels to promote TRPV1 sensitization^19–21^. Furthermore, some pro-algesic agents increase the exocytosis of vesicles carrying TRPV1, rising its expression on the neuronal surface^22,23^. Given the pivotal role of PLCβ in this inflammatory sensitization, peripheral PLCβ modulation is a valuable approach for the development of anti-inflammatory and antinociceptive drugs.

Currently, the aminosteroid U73122 ((1-(6-((17β-3-methoxyestra-1,3,5(10)-trien-17-yl)amino)hexyl)-1*H*-pyrrole-2,5-dione)) is widely used as a specific PLC inhibitor^24^. However, many studies have demonstrated the lack of specificity of this compound and off-target inhibitory effects^25–28^, partly due to its ability to alkylate cysteine groups^29^. Additionally, U73122 modulates TRP channels through a PLC-independent mechanism, being a potent agonist of TRPA1 and TRPM4 and a blocker of TRPM3^30,31^. Due to the off-target effects of U73122 and the involvement of PLCβ proteins in multiple signaling pathways, development of selective PLCβ inhibitors is crucial for understanding PLCβ-mediated cellular processes and for developing potential therapeutic agents.

The structure of PLCβ isoforms includes a N-terminal pleckstrin homology domain, four tandem EF hands, a catalytic triose phosphate isomerase (TIM) barrel domain, a C2 domain and an exclusive C-terminal region with a proximal and a distal domain^7,32–34^. Traditionally, the search of enzyme inhibitors was focused on compounds targeting the active site. Unfortunately, this protein region is well preserved among PLC isozymes and this complicates the identification of selective modulators. For instance, high-throughput screening pipelines identified three hit compounds that inhibited the activity of purified PLCβ3, but also of PLCδ1 and PLCγ1^35^. Intriguingly, nothing else is known about them. Another strategy for PLCβ inhibition was based on the necessary interaction with Gα_q_ proteins for PLCβ activation. Long peptides based on the helix-turn-helix motif in the proximal domain of PLCβ3, which contacts with Gα_q_, disrupted this protein-protein interaction^32,36^. However, through this site, Gα_q_ also interacts with other proteins such as p63RhoGEF, and these peptides also inhibited their activation. Altogether, these complications have prevented druggability of PLCβs.

In this scenario, two independent autoinhibitory mechanisms precisely regulate PLCβ isoforms to maintain a low basal activity^37^. The Hα2^’^ helix located in the proximal CT domain interacts with a region close to the active site and inhibits its enzymatic activity, until this Hα2^’^ helix is displaced by Gα_q_ binding^33^. Additionally, the XY linker connecting X and Y regions of the TIM barrel catalytic domain blocks the active site when PLCβ is inactive. This autoinhibitory loop is composed of an unconserved N-terminal region, followed by a high number of negatively-charged residues and an ordered region. This includes a short 3_10_ helix known as lid helix that occludes the active site^38^. Upon PLCβ activation, protein recruitment to the cell membrane promotes removal of the lid helix from the active center, driven by electrostatic repulsions between the acidic part of the linker and the negative charge on the membrane surface^37^. Thus, these autoinhibitory regions are unexploited targets for development of selective PLCβ modulators.

Here, we describe the discovery of small peptide analogues patterned after the XY linker as modulators of PLCβ enzymatic activity. We report that peptides derived from this autoinhibitory region bind to PLCβ isoforms and significantly inhibit their activity, as evidenced from PIP_2_ hydrolysis and intracellular calcium transients. Additionally, inhibition of the PLCβ pathway in a neuronal model attenuates bradykinin-induced action potential firing and TRPV1 sensitization. Finally, the best peptide in terms of activity and solubility shows promising anti-inflammatory and antinociceptive activity in a murine model of inflammatory pain. Collectively, our findings indicate that peptides patterned after the autoinhibitory XY linker act as selective PLCβ inhibitors, that can be tailored for other PLC isoforms to provide lead compounds for this family of undruggable targets.

## RESULTS

### PLCβ peptides patterned after the autoinhibitory XY linker

Using computational techniques of molecular modelling, we designed peptide analogues of the XY linker to inhibit the activity of PLCβ isoforms. As illustrated in **Figure 1A**, only a small region at the C-terminal end of the XY linker is crystallized in the PLCβ3 structural model (PDB code: 3OHM). This flexible region covers catalytic domain surface where the active center of PLCβ isoforms is located. In a first step, the XY linker was fragmented into smaller overlapping peptides sequentially moving from the N-terminus to the C-terminus with an offset of 1 amino acid (**Figure 1B**). Consequently, a set of shortened peptides containing 4 to 10 amino acids were selected from the XY linker using the structural context of PLCβ. This strategy allowed selected peptides to be bound to PLCβ3, while the remaining XY linker was removed. A total of 70 peptide-PLCβ3 complexes were modelled, and the interaction was graded by estimating the theoretical binding free energy (**Figure 1B** and **S1**). Normalization by peptide length suggests that small peptides (6-mer) exhibited values similar to longer peptides. Therefore, we focused on 6-mer peptides as they may exhibit more favorable physicochemical properties such as solubility, stability, and membrane permeability than longer counterparts.

**Figure 1.**
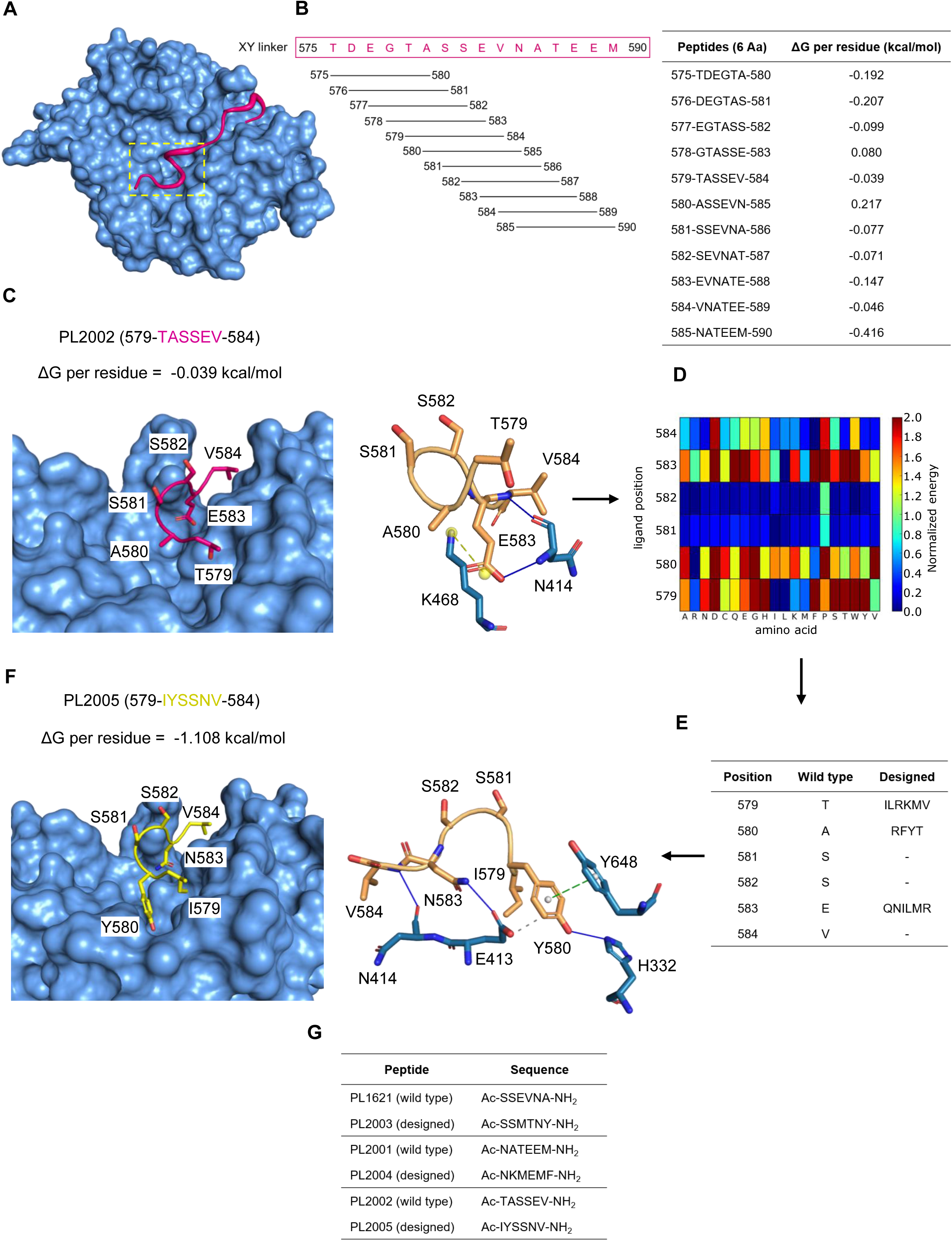
PLCβ peptides patterned after the autoinhibitory XY linker. **(A)** Catalytic domain of PLCβ3 isoform (blue surface) interacting with the crystallized region of autoinhibitory XY linker (pink cartoon). Approximate location of active site (dashed box). PDB code: 3OHM. **(B)** XY linker sequence cut into smaller overlapping peptides moving from the N-terminus to the C-terminus with an offset of 1. Predicted ΔG per residue of the resulting 6-mer peptides. More negative value means better interaction. **(C) Left panel**, anchor points of wild-type peptide 579-TASSEV-584 (pink, PL2002) bound to PLCβ3 isoform with predicted ΔG per residue. **Right panel**, PLCβ3 residues that directly interact with this peptide (blue). Solid blue lines: hydrogen bonds. Dashed yellow lines: salt bridges. **(D)** Sequence space search to optimize interaction of wild-type peptide with PLCβ3. Heat map shows normalized binding energies of the 20 natural amino acids (x-axis) at each position (y-axis) according to a color scale from blue (best fit) to red (worst). **(E)** The most favorable residues (Designed column) at each position (Position column) of the wild-type peptide (Wild type column) were combined to obtain new peptides. **(F) Left panel**, designed peptide 579-IYSSNV-584 (yellow, PL2005) shows increased theoretical affinity compared to wild-type (ΔG per residue). **Right panel**, PLCβ3 residues that directly interact with this peptide (blue). Solid blue lines: hydrogen bonds. Dashed green lines: π-stacking. Dashed grey lines: hydrophobic interactions. **(G)** Summary of PLCβ-modulating peptides selected for their functional validation. ΔG: binding free energy. Aa: amino acid. Ac: acetylated.

The 6-mer peptide with the most negative binding energy, and thereby the best interaction, was 585-NATEEM-590 (**Figure 1B**). This fragment comes from the C-terminus of the XY linker, distant from the active center. In this peptide, M590 is buried in the structure of PLCβ3, being a potential anchor point (**Figure S2A**). The other 6-mer peptides with the best binding energies are at the N-terminus of the XY linker (575-TDEGTA-580 and 576-DEGTAS-581). These fragments are derived from the XY region known as lid helix that physically blocks the active site of the protein^37^. In this region stands out residue E583, which is inserted into the binding interface, and it could be critical for the interaction with PLCβ3 (**Figure 1C**). Additionally, we focused on peptides having strong binding energies containing E583 (581-SSEVNA-586 and 579-TASSEV-584), together with peptide 585-NATEEM-590. Using this overlapping selection, the whole crystallized sequence of the XY linker was considered in the design strategy.

The binding of 6-mer peptides to the PLCβ3 catalytic center was evaluated in a sequence space search to obtain peptides with an improved theoretical binding free energy. Using a virtual mutagenesis approach, position-specific scoring matrices for each peptide were obtained with a prediction of the binding free energy of the 20 natural amino acids at each position (**Figure 1D**). Heat maps depict restrictive positions colored in red, where only specific residues fit, and permissive positions colored in blue, where the 20 amino acids show an optimal interaction. An example of a restrictive position in peptide 585-NATEEM-590 is M590 (**Figure S2B**). This residue is buried in a binding pocket, that only can accommodate the methionine or a hydrophobic amino acid. In contrast, S581 and S582 in peptide 579-TASSEV-584 are clear examples of permissive positions (**Figure 1D**). They are oriented towards the opposite direction of the PLCβ3 binding surface allowing almost any amino acid to be fitted. A selection of the best amino acids per position in terms of binding energy with different physicochemical properties were combined to obtain new sequences with an increased theoretical binding affinity, always keeping wild-type residues in permissive positions to save computational time (**Figure 1E**). Designed peptides obtained with their predicted binding energies can be found in the supplementary file Source_data.zip. Most of them show an improved binding energy compared to their corresponding wild-type sequence. For each wild-type peptide selected, a designed peptide ranked into the top 10 of binding energies, with a balance of polar and hydrophobic amino acids or positive and negative charges, was evaluated (**Figure 1G**).

Potential non-covalent interactions of wild-type and designed peptides to the PLCβ3 surface were predicted with the PLIP software. Peptide 579-TASSEV-584 (PL2002) interacts with PLCβ3 through residue E583, whose side chain forms a salt bridge with the amino group of the K468 side chain (**Figure 1C**), and creates a hydrogen bond with the N-group of the peptide bond between N414 and E413. Notably, the designed sequence 579-IYSSNV-584 (PL2005) improves its binding free energy (**Figure 1F**). The substitution of the alanine at position 580 by a tyrosine introduces a π-π interaction with the aromatic Y648 side chain. Besides, Y580 forms a hydrogen bond through its hydroxyl group with the imidazole of H332. The interchange of the critical E583 by an asparagine does not seem to affect the binding, creating a new hydrogen bond with the E413 side chain, whose γ-carbon also interacts with the Y580 aromatic ring.

The main anchor site for peptide 581-SSEVNA-586 (PL1621) is the residue E583, similarly to 579-TASSEV-584 (PL2002) (**Figure S3A**). In the designed peptide 581-SSMTNY-586 (PL2003), a new hydrogen bond is created through the interaction of the hydroxyl group of Y586 with the amino group of the R470 side chain (**Figure S3D**). Moreover, the aromatic ring of Y586 contacts with the β-carbon of K468. The mutant threonine introduced at position 584 establishes another hydrogen bond with D417 side chain and with the carbonyl group of the peptide bond between D417 and V416.

Finally, peptide 585-NATEEM-590 (PL2001) binds to PLCβ3 through the buried M590 and other contact points (**Figure S2A**). A586 establishes hydrophobic contacts through its side chains with the γ-carbon of R470 and the β-carbon of K468. The hydroxyl group of T587 forms a hydrogen bond with the guanidinium group of R470, while the β-carbon of its side chain contacts with the γ-carbon of Q422. Although E588 is not oriented to the binding surface, the deprotonated carboxyl group of its side chain at physiological pH could interact with the protonated guanidinium group of R470 through a salt bridge. Beyond, β and γ carbons of E589 interact with the side chain of A426 and A423, respectively. Interestingly, in the designed peptide 585-NKMEMF-590 (PL2004), phenylalanine introduction in the buried position 590 allows a T-shaped π-stacking interaction with the F412 aromatic side chain, and hydrophobic contacts with V465 and L593 (**Figure S2D**). Substitution of A586 by a lysine introduces a hydrogen bond with the carbonyl group of the peptide bond between K469 and R470.

In summary, six PLCβ peptides based on the autoinhibitory XY linker were computationally designed. These include three peptides with the wild-type XY sequence and their respective designed peptides with improved theoretical affinity by increasing the number of non-covalent interactions with the binding surface.

### PLCβ derived peptides inhibit PLCβ-mediated intracellular calcium transients

The inhibitory effect of PLCβ-modulating peptides was assessed recording the intracellular Ca^2+^ transients triggered after PLCβ activation by the muscarinic acetylcholine receptor 3 (m3AChR), endogenously expressed in HEK293 cells^39^. Stimulation with acetylcholine (ACh) elicited a robust calcium response (**Figure 2A**). Given the potential effect of ACh on mAChR and nicotinic receptors (nAChRs)^40,41^, ACh responses in HEK293 cells were characterized in Ca^2+^ microfluorimetry assays (**Figure S4**). Blockade of m3AChR with atropine abolished Ca^2+^ responses. Inhibition of the endoplasmic reticulum Ca^2+^-ATPase with thapsigargin (TG) also blocked ACh responses after depleting intracellular Ca^2+^ stores (**Figure S4A**). Furthermore, removal of Ca^2+^ from the extracellular solution did not affect ACh responses (**Figure S4B**). These results reveal that ACh responses in HEK293 cells are mediated by m3AChR coupled to PLCβ, discarding the potential involvement of endogenous nAChRs. This cell line also expresses the protein PLCβ3, used as a reference for the design of PLCβ-modulating peptides (**Figure S4C**).

**Figure 2.**
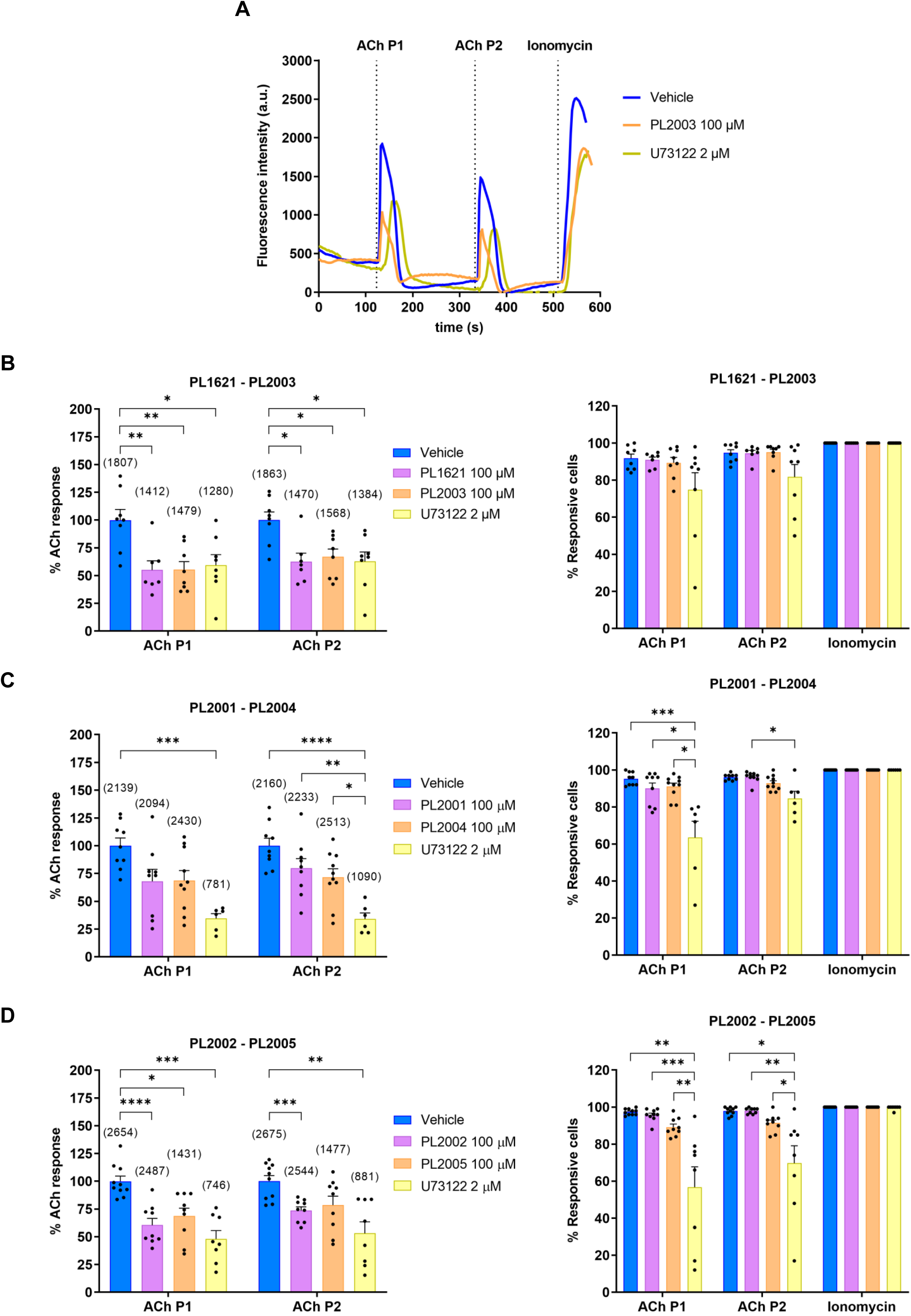
PLCβ-derived peptides inhibit PLCβ-mediated intracellular calcium transients. **(A)** Representative traces of intracellular Ca^2+^ transients to two pulses of 1 µM ACh (P1, P2) and a final pulse of 1 µM ionomycin in HEK293 cells treated with vehicle, one of the most potent PLCβ peptides (PL2003) or U73122. **(B-D) Left panels**, peptides patterned after the XY linker significantly attenuate ACh-induced intracellular Ca^2+^, except for PL2001-PL2004. The activity of wild-type XY sequence peptides (purple) is compared to that of designed peptides (orange) and U73122 (yellow). Percentage of ACh response normalized to vehicle. **Right panels,** PLCβ peptides do not significantly alter the percentage of ACh-responsive cells, unlike U73122. **(B-D)** Bar graphs indicate mean + Standard Error of Mean. Dots are the mean of each recording (N=3 biological replicates). Number of cells per condition in brackets in left panels. One-way ANOVA followed by Bonferroni or Kruskal-Wallis followed by Dunn’s test (*p<0.05, **p<0.01, ***p<0.001, ****p<0.0001). ACh: acetylcholine.

PLCβ activity was evaluated after applying two pulses of 1 µM ACh (**Figure 2A**). Peptides were tested in pairs to compare activity of wild type to mutated sequences. Peptides (100 µM) were pre-incubated for 1 h to facilitate cell permeability followed by target interaction. As illustrated **in Figures 2B-2D, left panels**, 6-mer peptides attenuated ACh-induced intracellular Ca^2+^ raise, showing a slight loss of inhibitory activity after the second stimulation. These peptides produced an inhibitory effect, similar to that of the unspecific PLC inhibitor, U73122. However, U73122 showed a high inter-assay variability in its inhibition efficacy **(Figure 2B-2D left panels)**.

As shown in **Figures 2B-2D, left panels**, there were not significant differences in the inhibitory potency of wild-type and designed peptides. PL1621 and PL2003 were the most potent peptides, inhibiting up to 45% of PLCβ activity in response to the first ACh pulse (44.9 ± 8.2% and 44.6 ± 7.2%, p<0.01 vehicle vs. PL1621 and PL2003, respectively, **Figure 2B, left panel**). The pair PL2001-PL2004 showed the best theoretical binding energies, but they caused a lower and non-significant reduction of intracellular Ca^2+^ levels compared to PL1621 and PL2003 (32.0 ± 10.7% and 31.3 ± 8.9% in the presence of PL2001 and PL2004, respectively, **Figure 2C, left panel**). PL2002 also induced a significant reduction of ACh responses (39.5 ± 5.9%, p<0.0001 vs. vehicle, **Figure 2D, left panel**). Unfortunately, its designed counterpart PL2005 showed solubility problems in different solvents. Nevertheless, a modest inhibitory activity was observed (31.3 ± 7.1%, p<0.05 vs. vehicle, **Figure 2D, left panel**). In most assays, U73122 significantly reduced the number of cells responding to ACh without compromising ionomycin responses (**Figures 2C and 2D, right panels**). Unlike U73122, PLCβ-modulating peptides did not significantly alter the percentage of ACh responsive cells (**Figures 2B-2D, right panels**). Together, these data suggest that peptides patterned after the XY linker induce a partial catalytic inhibition of PLCβ enzymes.

### PL2003 inhibits PIP_2_ hydrolysis in cells

To further investigate the mode of action of designed inhibitory PLCβ peptides, we selected PL2003 as representative for evaluating its effect on the hydrolysis of membrane PIP_2_. The level of PIP_2_ was optically monitored with the fluorescent construct PLCδ-PH-GFP, encoding a selective PIP_2_ binding domain (PH) conjugated to the green fluorescent protein (GFP)^42^. For this purpose, HEK293T cells were co-transfected with the PIP_2_ reporter and the bradykinin receptor 2 (B_2_R) that also signals through PLCβ. Under basal conditions, the fluorescent reporter is located to the inner leaflet of the plasma membrane bilayer, interacting with PIP_2_, with minimal expression in the cytosol (**Figure 3A, left panels**). In contrast, bradykinin (BK) stimulation of B_2_R activates PLCβ enzymatic activity, which hydrolyses PIP_2_, promoting the translocation of the reporter PLCδ-PH-GFP to the cytosol (**Figure 3A, right panels**) (videos in supplementary materials). Translocation of PLCδ-PH-GFP caused a remarkable increment of cytosolic fluorescence in HEK293 cells pre-incubated with vehicle (**Figure 3B**). Noteworthy, 100 µM peptide PL2003 pre-incubated for 1 h, attenuated by 40% the cytosolic fluorescence increase induced by BK (**Figures 3B and 3C**), akin to 2 µM U73122 (0.127 ± 0.009 and 0.126 ± 0.009, p<0.0001 vs. vehicle in the presence of PL2003 and U73122, respectively). In contrast, neither PL2003 nor U73122 modified the time to reach the maximum PLCβ activity, measured from the onset of PLCδ-PH-GFP translocation (**Figure 3D**). Normalization of representative PLCδ-PH-GFP traces to their maximum and minimum values yielded traces with similar slopes (**Figure 3E**). These findings indicate that PL2003 does not alter the kinetics of PIP_2_ hydrolysis, exhibiting a reaction velocity almost identical to that reported in the presence of vehicle.

**Figure 3.**
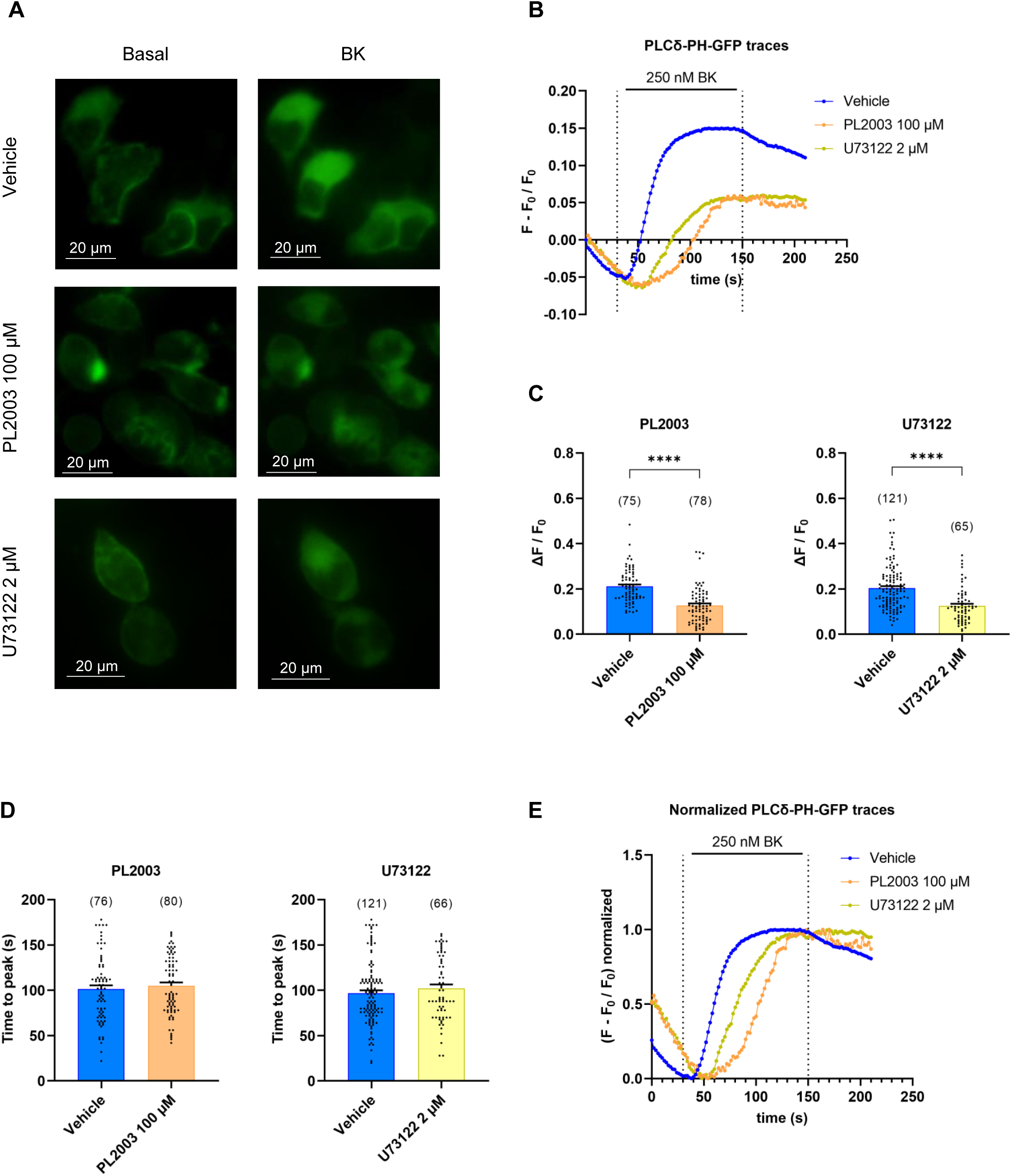
PL2003 inhibits PIP_2_ hydrolysis. **(A)** Representative pictures of HEK293T cells transfected with the PIP_2_ probe PLCδ-PH-GFP and pre-treated with vehicle, PL2003 or U73122, before (basal) and after applying bradykinin (BK). **(B)** Representative fluorescence traces illustrating BK-induced translocation of PLCδ-PH-GFP from plasma membrane to cytosol. A slight fluorescence rundown is observed during basal fluorescence assessment. PL2003 attenuates PLCδ-PH-GFP translocation similarly to PLC inhibitor U73122. **(D)** PL2003 or U73122 does not modify the time to reach maximum translocation. **(E)** The slope of PIP_2_ hydrolysis is similar after vehicle, PL2003 or U73122. **(C,D)** Data expressed as mean + Standard Error of Mean. Dots on bar charts are cells from N=3 or 4 biological replicates in presence of PL2003 or U73122, respectively. Number of cells analyzed in brackets. Mann-Whitney U (****p<0.0001). BK: bradykinin.

These data suggest that PL2003 directly inhibited the enzymatic activity of PLCβ isoforms reducing the hydrolysis of the membrane PIP_2_ and the release of IP_3_-mediated Ca^2+^ from the endoplasmic reticulum.

### Positively-charged amino acids at the N-terminus of PL2005 improve solubility preserving activity

PL2003 was formulated in saline solution for later *in vivo* testing, but a strong aggregation was observed conditioning its potential therapeutic testing. Thus, we first designed an aqueous soluble 6-mer peptide for *in vivo* pharmacological studies. The designed peptide PL2005 (579-IYSSNV-584), exhibited one of the best binding free energies (**Figure 1F**). However, this peptide was initially discarded due to its moderate potency blocking ACh responses (31.3 ± 7.1%, p<0.05, **Figure 2D, left panel**) and its poor aqueous solubility. Nonetheless, because its apparent high binding energy we reasoned that this peptide could be a good therapeutic candidate and it was selected for improving its solubility. For this task, two positive residues (KR) were introduced at the N-terminus to improve water solubility and cell penetration, particularly of neuronal membranes that exhibit a negative resting membrane potential. As depicted in **Figure 4A**, the addition of K577 and R578 notably improved the binding energy when compared to the original sequence (577-EGIYSSNV-584). Peptide 577-KRIYSSNV-584 (hereinafter named PL2204) showed a normalized binding free energy similar to PL2005 (ΔG per residue: −1.108 kcal/mol). The guanidinium group of R578 interacts with the Q340 side chain through hydrogen bonds, creating a new anchor point (**Figure 4B**). Moreover, the bulky and polar side chain of R578 rotates the side chain of the adjacent I579, which contacts with the β-carbon of the D364 side chain and the indole W366.

**Figure 4.**
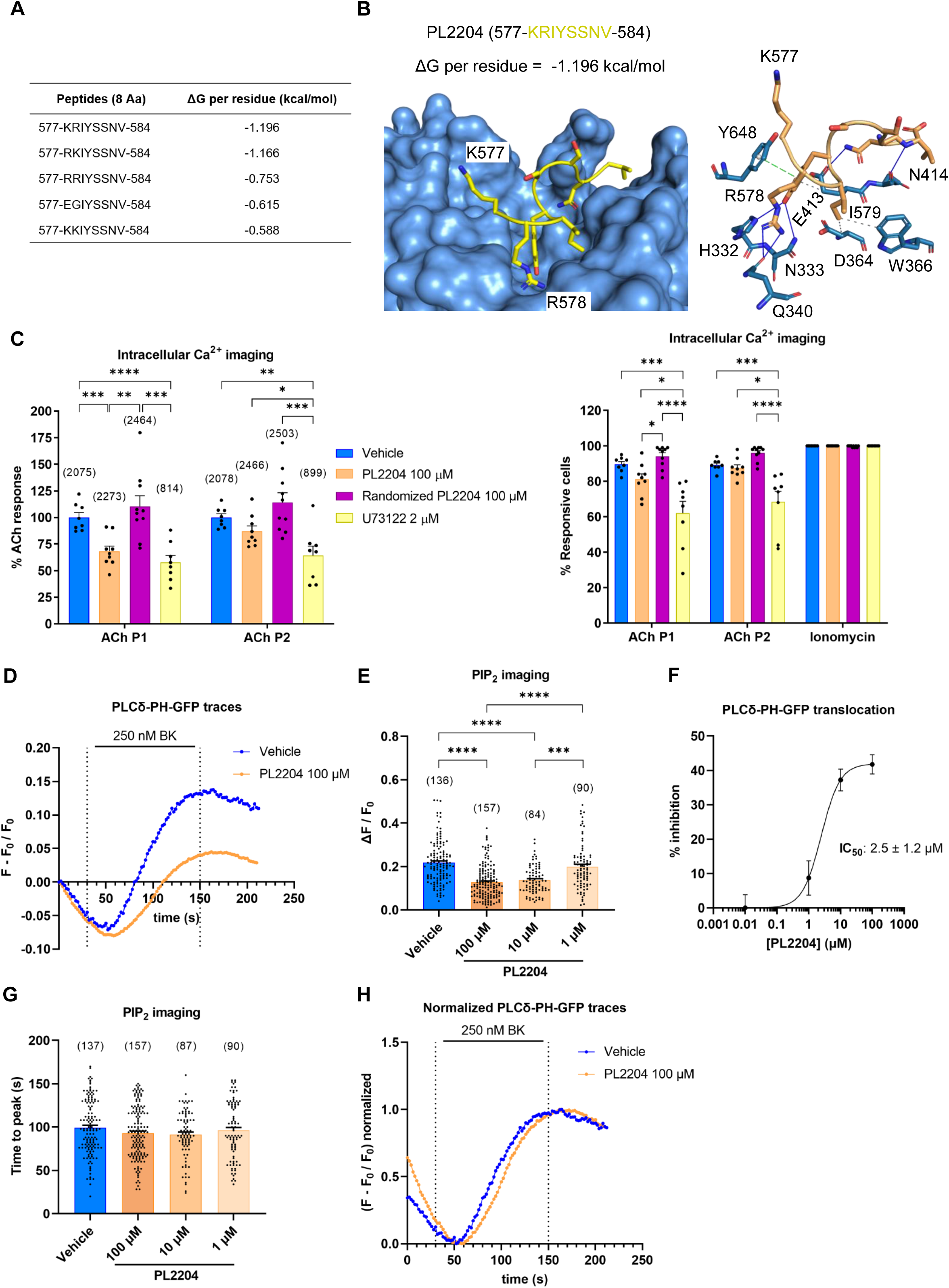
Positively-charged amino acids at the N-terminus of PL2005 improve solubility while preserving activity. **(A)** ΔG per residue of 8-mer peptides obtained by addition of two positive charges at the N-terminus of PL2005 (579-IYSSNV-584). Peptide 577-EGIYSSNV-584 containing wild-type residues at positions 577 and 578 included. More negative value means better interaction. **(B) Left panel,** designed 8-mer peptide 577-KRIYSSNV-584 (yellow, PL2204) shows similar ΔG to the original 6-mer PL2005. **Right panel,** interactions between the 8-mer peptide and PLCβ3. PLCβ3 residues directly interacting in blue. Solid blue lines: hydrogen bonds. Dashed green lines: π-stacking. Dashed grey lines: hydrophobic interactions. **(C) Left panel,** PL2204 attenuates ACh-induced intracellular Ca^2+^ transients, while a randomized PL2204 peptide not. Percentage of ACh response normalized to vehicle. **Right panel,** percentage of ACh-responsive cells is similar after PL2204 or vehicle. **(D)** Representative fluorescence traces illustrating BK-induced translocation of PLCδ-PH-GFP from plasma membrane to cytosol in vehicle or PL2204 (100 µM). A slight fluorescence rundown is observed during basal fluorescence assessment. **(E)** 10-100 µM PL2204 attenuates PLCδ-PH-GFP translocation-associated fluorescence. **(F)** Concentration-dependent inhibitory effect of PL2204. IC_50_ value of 2.5 ± 1.2 µM (mean ± SEM). **(G)** Similar time to reach maximum translocation after 1-100 µM PL2204 or vehicle. **(H)** The slope of PIP_2_ hydrolysis is similar in PL2204 or vehicle. **(C,E,G)** Data of bar charts expressed as mean + Standard Error of Mean. **(C)** Dots are the mean of each recording from N=3 biological replicates. Numb er of cells per condition in brackets. **(E,G)** Dots are cells N≥3 biological replicates. **(C,E,G)** One-way ANOVA followed by Bonferroni or Kruskal-Wallis followed by Dunn’s test (*p<0.05, **p<0.01, ***p<0.001, ****p<0.0001). ΔG: binding free energy. Aa: amino acid. ACh: acetylcholine. BK: bradykinin.

PL2204 was soluble in a physiological saline solution and DMSO. The activity of PL2204 on the intracellular Ca^2+^ transients triggered by ACh was tested following the previously described protocol. PL2204 at 100 µM induced a reduction of ACh responses of approximately 30% (31.8 ± 4.9%, p<0.001 vs. vehicle), revealing that modifications in the sequence of PL2005 did not change its PLCβ inhibitory activity (**Figure 4C, left panel**). The sequence dependence of its inhibitory activity was validated using a randomized peptide containing the same amino acids composition in a random sequence. The randomized PL2204 completely lost its inhibitory activity, indicating that PLCβ inhibition depends on the peptide amino acid sequence. As with previous PLCβ peptides, PL2204 did not significantly reduce the percentage of ACh-responsive cells when compared to vehicle-treated cells (**Figure 4C, right panel**).

PL2204 also inhibited the hydrolysis of PIP_2_ and attenuated the BK-triggered translocation of the PIP_2_ probe PLCδ-PH-GFP from the cell membrane to the cytosol (**Figures 4D and 4E**) (videos in supplementary materials). As shown in **Figure 4F**, PL2204 followed a sigmoidal dose-response relationship in the micromolar range with a maximum inhibition at 100 µM (41.8 ± 2.8%, p<0.0001 vs. vehicle). The peptide induced similar inhibition at 10 µM (37.2 ± 3.2%, p<0.0001 vs. vehicle), whereas it has almost no effect at 1 µM (8.7 ± 5.0%). The IC_50_ was 2.5 ± 1.2 µM. Like PL2003 and U73122, PL2204 did not alter the time required to reach the maximum PLCδ-PH-GFP translocation (**Figure 4G**), displaying identical kinetics to vehicle (**Figure 4H**).

Taken together, these results support that rational addition of two positive charges to PL2005 resulted in a more soluble peptide in saline solutions. Although PL2003 appears to be more potent than PL2204 in calcium imaging screenings (44.7 ± 7.1% vs. 31.8 ± 4.9% inhibition with PL2003 and PL2204, respectively), both peptides similarly inhibit the PLCβ activity in the PIP_2_ assay (39.8 ± 4.2% vs. 41.8 ± 2.8% with PL2003 and PL2204, respectively), suggesting a similar *in vitro* potency.

### PLCγ activity is preserved in the presence of PL2003 and PL2204

The selectivity of PLCβ-modulating peptides for PLCβ isoforms was assessed recording Ca^2+^ transients evoked by store-operated calcium channels (SOC). Previous works demonstrated that SOC activation in human keratinocytes requires PLCγ activity^43^. To evaluate this mechanism, HaCaT cells were treated with thapsigargin in Ca^2+^-free extracellular solution. TG depleted intracellular Ca^2+^ stores, thereby triggering SOC activation through a PLCγ-dependent mechanism. Subsequent addition of extracellular Ca^2+^ elicited a robust response as a result of Ca^2+^ influx through SOC (**Figure 5A**). This process was significantly inhibited in cells pre-treated with the unspecific PLC inhibitor U73122 (**Figures 5A and 5B, left panels**). In contrast, SOC activity remained unaltered in the presence of PL2003 or PL2204 compared to vehicle-treated cells (**Figure 5A, right panel, and 5B, middle and right panels**). These findings suggest that these PLCβ-modulating peptides did not alter PLCγ activity, thus substantiating their selectivity for PLCβ isoforms.

**Figure 5.**
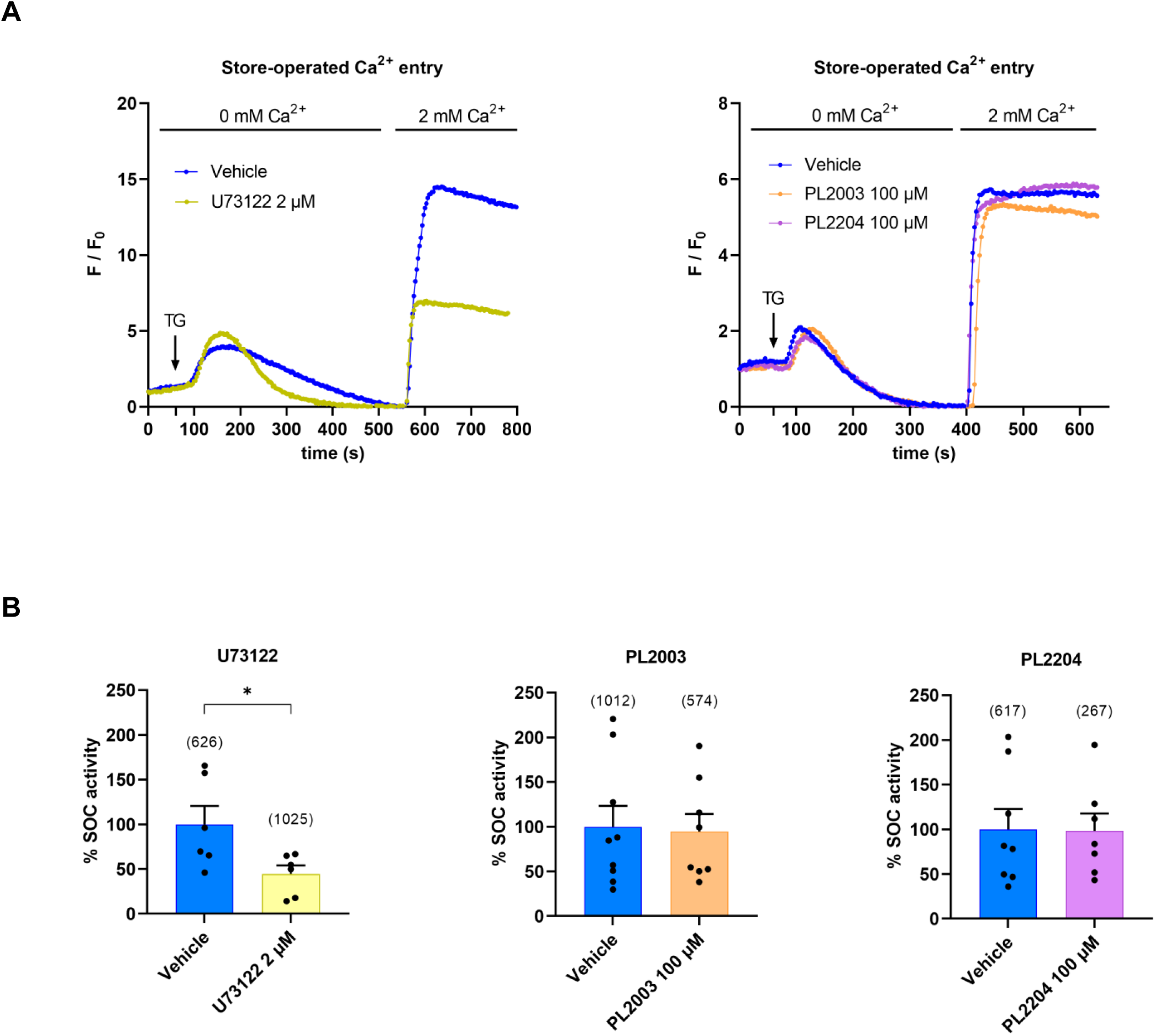
PLCγ activity is preserved in the presence of PL2003 and PL2204. **(A)** Representative fluorescence traces of TG-induced SOC activity in HaCaT cells treated with vehicle, U73122 (left panel) or PL2003 and PL2204 (right panel). Depletion of intracellular calcium stores by 1 µM TG in Ca^2+^-free extracellular solution activates SOC through a mechanism partly dependent on PLCγ. Re-addition of extracellular calcium triggers SOC-mediated calcium influx. **(B)** U73122 attenuates SOC response, whereas PL2003 and PL2204 have no effect. SOC activity expressed as the ratio ΔF_SOC_ / ΔF_TG_, normalized to vehicle-treated cells. **(B)** Data expressed as mean + Standard Error of Mean. Dots are the mean of each recording (N=3 or 2 biological replicates for PL2003-PL2204 or U73122, respectively. Number of cells per condition in brackets. Unpaired t-test (*p<0.05). TG: thapsigargin. SOC: store-operated calcium channels.

### PL2003 and PL2204 inhibit BK-induced neuronal electrogenic activity and TRPV1 sensitization

BK induces PIP_2_ hydrolysis through PLCβ, leading to PKC activation by DAG and Ca^2+^ that, in turn, phosphorylates TRPV1 potentiating its activity^9,21^. In addition, BK promotes neuronal excitability by modulating the activity of other ion channels^44^. Thus, we next investigated whether PLCβ inhibition could attenuate BK-mediated neuronal electrogenesis and TRPV1 sensitization in sensory neurons. For this purpose, primary cultures of neonatal rat dorsal root ganglion (DRG) neurons were seeded on multielectrode arrays (MEA) to monitor neuronal networks activity. Repeated neuronal stimulation with three 500 nM capsaicin pulses (P1, P2 and P3), interspersed with washing periods, induced a progressive TRPV1 desensitization, evidenced by a significant decrease in the neuronal firing evoked by P2 and P3 (**Figures 6A and 6C**). However, application of 1 µM BK between P2 and P3 for 8 min promoted an increase in the neuronal electrogenic activity and prevented desensitization of P3 (**Figure 6B first panel and 6D**). These data indicate that BK activation of PLCβ signaling hyperexcites sensory neurons incrementing their electrical firing.

**Figure 6.**
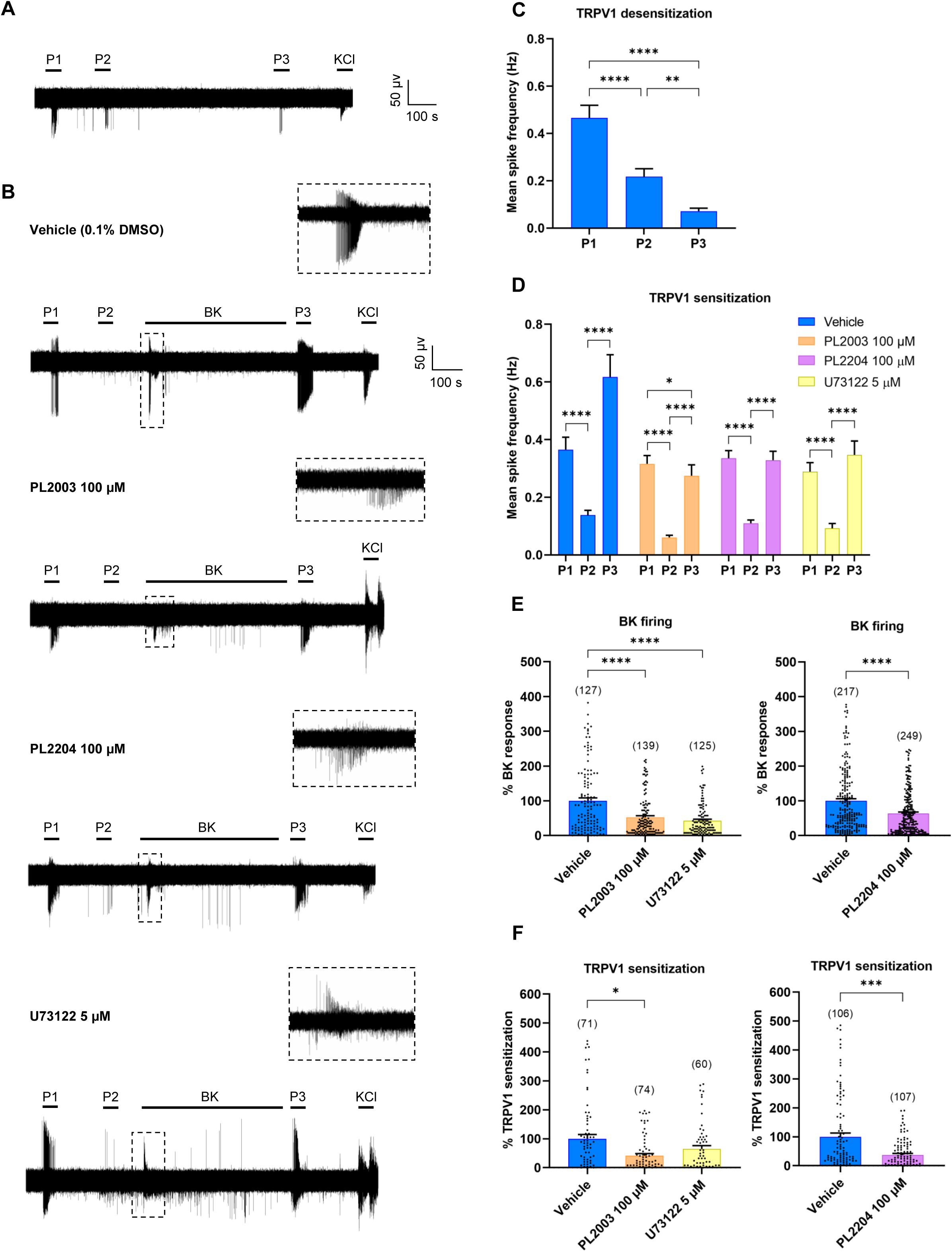
PL2003 and PL2204 inhibit BK-induced neuronal electrogenic activity and TRPV1 sensitization. Representative recordings of TRPV1 activity in DRG neuronal cultures **(A)** non-sensitized and **(B)** sensitized with BK in the presence of PL2003 or PL2204, its vehicle or PLC inhibitor U73122. Three pulses of 500 nM capsaicin (P1, P2, P3) were applied. In the desensitization protocol **(A)**, extracellular solution was applied between P2 and P3. In sensitization protocols **(B)**, 1 µM BK was applied to activate the PLCβ pathway. Magnified view of BK-induced action potentials within dashed squares. **(C, D)** Mean spike frequency after each pulse of capsaicin (P1, P2, P3) in non-sensitized **(C)** and BK-sensitized cultures. **(E)** PL2003 and PL2204 significantly reduce BK-induced action potentials. Mean spike frequency normalized to vehicle. **(F)** PL2003 and PL2204 attenuate BK-induced TRPV1 sensitization, expressed as P3 – P2 response normalized to vehicle. **(C-F)** Data expressed as mean + Standard Error of Mean. **(E,F)** Dots on bar charts are data from recording electrodes from N=3 biological replicates. Number of electrodes recorded in brackets. **(C-F)** Kruskal-Wallis followed by Dunn’s test (*p<0.05, **p<0.01, ****p<0.0001) or Mann-Whitney U (***p< 0.001, ****p<0.0001). BK: bradykinin.

We hypothesized that our best PLCβ-modulating peptides may attenuate both the BK-induced TRPV1 sensitization as well as its direct electrogenic activity. Neuronal cultures were incubated for 1 h with PL2003 or PL2204 (100 µM), its vehicle (0.1% DMSO) or the U73122. As depicted in **Figure 6E left panel**, PL2003 significantly reduced the firing of BK-evoked action potentials by nearly 50% compared to vehicle (47.7 ± 4.8% inhibition, p<0.0001), reproducing U73122 inhibitory effect (57.1 ± 4.0% inhibition, p<0.0001). Noteworthy, PL2003-mediated PLCβ inhibition also attenuated BK-induced TRPV1 potentiation compared to vehicle (**Figure 6F, left panel**). The presence of PL2003 resulted in a significant reduction of capsaicin evoked action potentials (P3) as compared to vehicle (58.2 ± 7.2% inhibition, p<0.05, **Figure 6F, left panel**). Intriguingly, although U73122 was more effective than PL2003 reducing BK-mediated action potential firing, it did not attenuate BK-induced TRPV1 sensitization (**Figure 6F, left panel**).

PL2204 also caused a significant reduction in BK-evoked action potential firing compared to vehicle (**Figure 6E, right panel**, 36.1 ± 3.9% inhibition, p<0.0001). Furthermore, as shown in **Figure 6F right panel**, PL2204 reduced TRPV1 sensitization by 60% compared to vehicle (62.6 ± 5.1% inhibition, p<0.001). The improved PL2204 peptide inhibited BK-induced electrogenicity and TRPV1 sensitization similarly to PL2003. Therefore, due to its enhanced solubility in saline, PL2204 was selected for *in vivo* evaluation.

### PL2204 displays *in vivo* anti-inflammatory and antinociceptive activity

Given the prominent *in vitro* effects of PL2204, we conducted an *in vivo* behavioral experiment in mice to investigate a possible effect of the peptide in preventing inflammation and nociceptive hypersensitivity. Mice received an intraplantar dose of PL2204 (250 µg) or vehicle, and 2 h later an injection of CFA or saline at the same site. This yielded 3 groups: (1) CFA-injected mice treated with PL2204, (2) CFA-injected mice treated with vehicle, and (3) a saline-injected group treated with vehicle (**Figure S5**). Inflammation was measured 2 h after CFA injection, whereas mechanical and heat hypersensitivity were assessed 3 and 4 h post-injection, respectively.

Mice treated with vehicle and injected with CFA developed strong and persistent inflammation for 7 days (**Figure 7A**, p<0.001 vs. baseline and control group Vehicle+Saline). In contrast, PL2204-treated mice revealed a reduced inflammation 2 h and 1 d after CFA (**Figure 7A**, p<0.001 vs. Vehicle+CFA). This anti-inflammatory effect gradually disappeared at 96 h, when swelling of ipsilateral paw became similar between vehicle and PL2204 treated mice (**Figure 7A**, 4-7 d).

**Figure 7.**
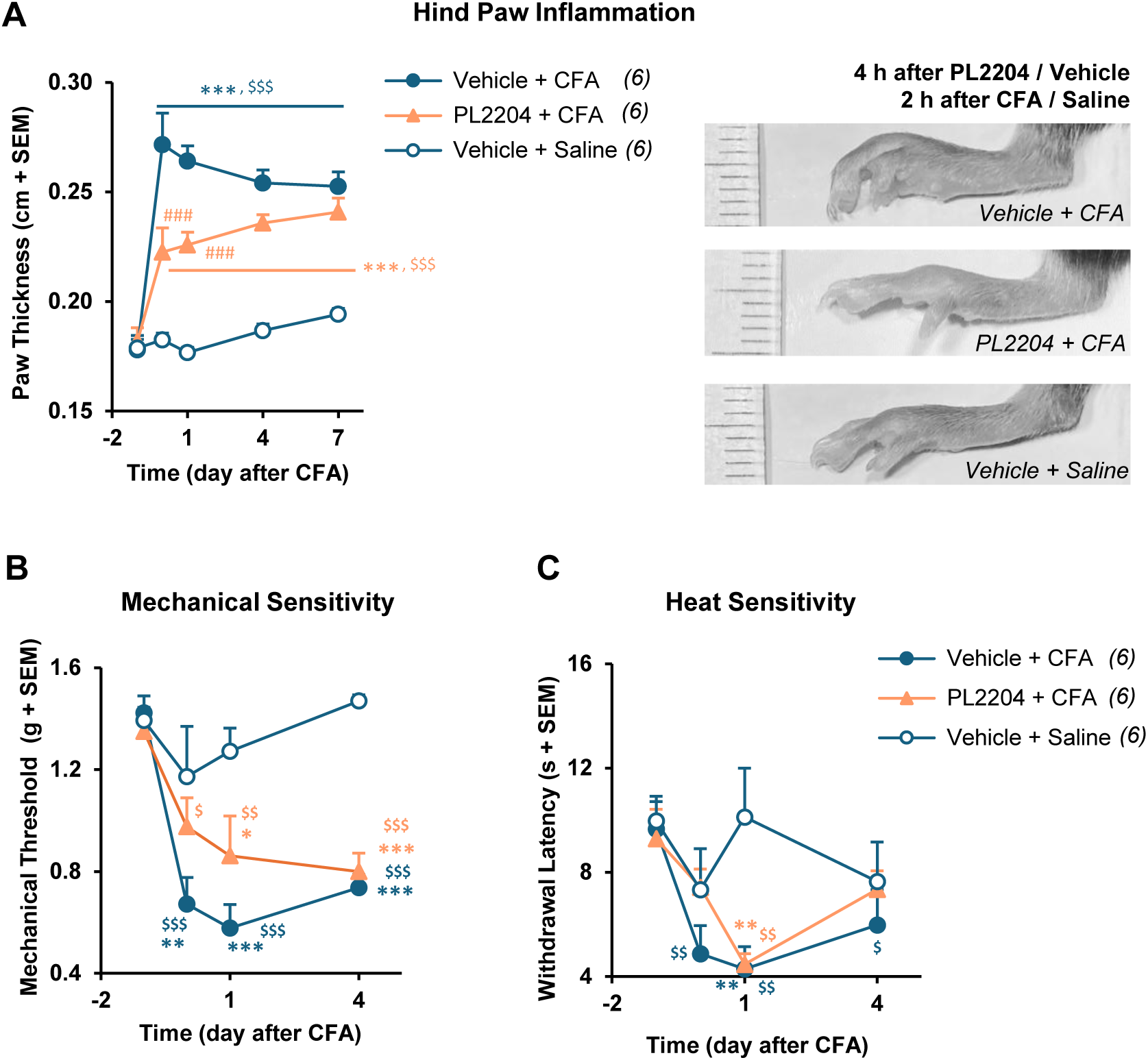
PL2204 displays *in vivo* anti-inflammatory and antinociceptive activity. **(A) Left panel,** mice treated with PL2204 showed less CFA-induced inflammation than those treated with vehicle and injected with CFA. **Right panel**, representative pictures showing reduced CFA-induced inflammation after pre-treatment with PL2204 or vehicle **(B)** Mice treated with PL2204 showed alleviation of mechanical hypersensitivity 3 h after CFA (N.S. vs. Vehicle+Saline). On the contrary, CFA-injected mice treated with vehicle developed robust mechanical hypersensitivity at all time points. **(C)** Mice treated with PL2204 showed slightly reduced heat sensitization 4 h after CFA (N.S. vs. Vehicle+Saline, N.S. vs. baseline) whereas CFA-injected mice treated with vehicle showed significant heat hypersensitivity at 4 h and 1 d. **(A-C)** Mean + Standard Error of Mean (SEM) values are shown. Two-way Anova and mixed models followed by Tukey and Dunnett’s tests. *p<0.05, **p<0.01, ***p<0.001 vs. control (Vehicle+Saline). ^$^p<0.05, ^$$^p<0.01, ^$$$^p<0.001 vs. baseline. ^###^p<0.001 vs. Vehicle+CFA. N=6 animals per group. CFA: complete Freund’s adjuvant.

CFA also induced pronounced mechanical hypersensitivity in vehicle-treated mice, significant from 3 h to 4 d after injection (**Figure 7B**, p<0.001 vs. baseline and p<0.01-0.001 vs. control group). On the contrary, PL2204-treated mice showed significant alleviation of mechanosensitivity 3 h after CFA (**Figure 7B**, N.S. vs. control group). Although, hypersensitivity emerged 24 h later in this group (**Figure 7B**, p<0.05 vs. control group and p<0.01 vs. baseline). In terms of heat nociception, vehicle-treated mice developed significant hyperalgesia 4 h after CFA (**Figure 7C**, p<0.01 vs. baseline) which was slightly reduced in PL2204 pre-treated mice.

After investigating the preventive effects of PL2204, its therapeutic activity was assessed. Eight days after CFA, when all animals with inflammation reached similar paw thickness and mechanical hypersensitivity, mice that were initially treated with vehicle were exposed for the first time to PL2204, whereas mice previously exposed to PL2204 were now treated with vehicle. The control group was kept without inflammation and received also vehicle. Two hours after the treatments, mice newly exposed to PL2204 showed a slight reduction in paw thickness when compared to previous measurements (**Figure S6A**, p<0.05 vs. day 7) and showed a partial antinociceptive effect (**Figure S6B,** N.S. vs. Vehicle+Saline+Vehicle or PL2204+CFA+Vehicle). On the contrary, vehicle-treated mice showed unchanged paw swelling and mechanical hypersensitivity (**Figure S6**). Hence, therapeutic activity of PL2204 could still be observed at this later stage of the inflammatory process, although such activity was less pronounced than when administered as a preventive treatment.

## DISCUSSION

The salient contribution of this study is the innovative development of peptide inhibitors targeting PLCβ enzymes through the combination of *in silico*, *in vitro* and *in vivo* complementary approaches. We report that small peptides derived from the autoinhibitory XY linker of PLCβ3 reduced PIP_2_ hydrolysis by inhibiting PLCβ enzymatic activity, resulting in an attenuation of inflammation-induced neuronal electrogenesis and TRPV1 sensitization. These peptides are sequence- and isoform-specific, since a random sequence peptide was inactive and peptides after the PLCβ XY linker did not affect PLCγ activity. PL2204, an aqueous soluble 8-mer peptide incorporating two positively-charged residues at the N-terminus, exhibited *in vivo* anti-inflammatory and antinociceptive activity in the CFA mouse model of inflammatory pain. These peptides represent a significant progress towards the development of novel pharmacological leads and enable druggability of these enzymes that have notably resisted pharmacological targeting.

We hypothesized that the autoinhibitory XY linker, a region that physically occludes the catalytic site of PLC enzymes, could be a source of peptide-based inhibitors and used a rational design approach to identify candidates within this motif. XY linkers with different amino acid sequence can be found in PLCβ, PLCδ and PLCε isoforms, while PLCγ isoforms have a more complex XY region with multiple domains^38,45^. The sequence diversity of this protein region within different PLC isoforms may facilitate the development of isoform-selective inhibitors, a yet unmet goal for this enzyme family. Here, we focused on PLCβ taking advantage of the availability of the PLCβ3 structure. Analysis of the XY linker sequence reveals that most of the residues are conserved (62.5%) among PLCβ1-β2-β3 isoforms, and more divergent when compared to PLCβ4 (**Figure S7A**). In contrast, PLCβ3 residues harboring the XY binding site are mostly conserved among the four PLCβ isoforms (**Figure S7B**), suggesting potential cross-reactivity of peptides patterned after the XY linker. Hence, it is plausible that our small peptides primarily target PLCβ3 but may cross-interact to some extend with the other PLCβ isoforms. Nonetheless, sequence differences in the XY linker provide the basis to rationally evolve isoform-selective sequences for this enzyme family.

The activity of PLCβ-modulating peptides was assessed *in vitro* using m3AChR and B_2_R receptors coupled to PLCβ isoforms. PLCβ peptides patterned after the XY linker attenuated ER-Ca^2+^ release triggered by m3AChR activation, while maintaining the number of responsive cells. Interestingly, the amplitude of Ca^2+^ transients elicited after a second ACh pulse was still partially reduced after the washout period. These findings are compatible with the mode of action of a moderate-affinity PLCβ inhibitor that hinders PIP_2_ access to the catalytic site, partially inhibiting enzymatic activity. Unlike glue-type drugs, these molecules with moderate potency may have the therapeutic advantage of efficiently targeting pathological states while preserving the activity of physiologically working enzyme pools. Such a partial inhibition of PLCβ was corroborated with the most potent peptide PL2003 and PL2204 in a PIP_2_ hydrolyzing assay, revealing direct effect on the hydrolysis of the membrane phospholipid PIP_2_. In addition, these peptides did not slow the kinetics of PIP_2_ hydrolysis, consistent with a competitive inhibitory mechanism between PL2003 or PL2204 and PIP_2_ for the active site of PLCβ. Furthermore, PL2003 and PL2204 did not affect store-operated calcium entry in HaCaT cells, a process dependent on PLCγ activity, which underscored their selectivity for PLCβ. This selectivity distinguishes our peptides from the unspecific PLC inhibitor U73122, thus allowing specific modulation of PLCβ-dependent mechanisms while preserving the activity of other relevant PLC isoforms such as PLCγ.

The moderate potency and efficacy of peptides may be influenced by several factors related to the cellular assay used. First, access to the enzyme is restricted by the cell membrane as the peptide-binding site is located intracellularly. Thus, peptide permeability appears pivotal for potent inhibition. Second, since PLCs are membrane proteins that hydrolyze lipids, their catalytic site may be hindered in the internal membrane leaflet, limiting access to our peptides. In addition, the accessibility of the catalytic site is limited in basal conditions as it is occupied by the XY linker and only accessible when the enzyme is activated. Thus, designed peptides must compete with the enzyme’s own XY linker for binding. Third, small peptides made of natural L-amino acids may be metabolized by cytosolic proteases, thus reducing their effective concentration for enzyme inhibition. These factors may account for the relatively high concentration of peptide (10-100 µM) required for PLCβ inhibition. Nevertheless, considering these limitations, it is remarkable that 6/8-mer peptides exhibited micromolar inhibitory activity.

Strategies such as the use of cell penetrating peptides or lipidation have been used to increase membrane permeability and anchorage of peptides. The most frequently used cell-penetrating peptide is the Tat sequence, which is rich in positively charged amino acids and takes advantage of the negative membrane potential to enhance membrane permeability^46^. This strategy is very efficient in translocating peptides, although due to its high positive charge tends to drive peptides to the cell nucleus which exhibits strong negative charge. We used this approach in a conservative manner to increase the solubility of PL2005 by incorporating two positively charged amino acids at the N-terminus, mimicking naturally occurring cell-penetrating peptides, and to minimize nuclear localization^46^. With this modification, our 8-mer peptide PL2204, with a net positive charge (+2), inhibited PLCβ enzymatic activity. Notably, a randomized version of PL2204 lost the inhibitory activity, underscoring the relevance of the specific order of amino acids for the interaction with the binding surface of PLCβ.

Peptide lipidation is a strategy that anchors peptide sequences into the lipid bilayer increasing their membrane concentration^47^. This modification has been used to modulate protein-protein interactions at the membrane surface such those involved in the SNARE complex^48^, the interaction of GPCRs with the heterotrimeric proteins^49^ and the interaction of intracellular protein domains involved in channel gating^50^. This strategy appears less appropriate to favor the binding of peptides to a catalytic site that most likely requires flexibility of the inhibitor. However, its degrees of freedom are limited by the rigid anchorage to the internal lipid monolayer imposed by the lipid. To circumvent this limitation, the use of flexible linkers for proper mobility is required. Although we do not fully discard this strategy to enhance the potency of our peptides, we do not prioritize it because peptide lipidation often increase peptide toxicity most likely by affecting the membrane physicochemical properties that may affect signaling processes. An alternative strategy is represented by the encapsulation of peptides in liposomes that also favor membrane permeability.

Our peptides also significantly attenuated BK-induced electrogenicity in sensory neurons, consistent with a PLCβ-dependent mechanism of BK-induced excitability. Our findings agree with growing evidence suggesting that depolarizing effects of bradykinin in nociceptive neurons are mediated by inhibition of M-type K^+^ channels and opening of Ca^2+^-activated chloride channels such as ANO1^44^. These events depend on the rise of intracellular calcium triggered by the PLCβ-IP_3_ pathway^44,51^. PLCβ signaling is also involved in inflammatory sensitization of TRPV1 channels expressed in sensory neurons^52–54^. Indeed, several algogens (ATP, BK, histamine, etc.) potentiate TRPV1 activity by increasing IP_3_ and DAG through activation of neuronal PLCβ. Notably, our peptides reduced BK-elicited TRPV1 sensitization. Inhibition of PLCβ signaling with our peptides was less potent in preventing TRPV1 inflammatory potentiation than direct B_2_R blockade with the specific antagonist HOE140^22^. This pharmacological approach likely causes stronger inhibition because B_2_R also activate Gα_s_ proteins coupled to adenylate cyclase^55^. Specifically, adenylate cyclase generates cAMP that activates protein kinase A, which also phosphorylates and sensitizes TRPV1^56^. Since PLCβ isoforms are not involved in this pathway, a degree of TRPV1 potentiation is expected after the PLCβ-directed peptides.

While our main focus was the study of BK response, the inflammatory soup contains additional mediators that trigger PLCβ activation, including histamine, protease-activated receptor 2 or ATP^14,52–54^. The effect of these mediators should then be also sensitive to the PLCβ-modulating peptides, and a stronger overall effect on inflammation could be expected. Furthermore, the PLCβ pathway modulates other members of the TRP family under inflammatory conditions^9^. For instance, TRPA1, a sensor of environmental irritants and endogenous proalgesic agents involved in inflammatory pain^57^, undergoes sensitization through PLCβ-dependent mechanisms^58,59^. Thus, inhibition of PLCβ should attenuate the sensitization triggered by multiple pro-nociceptive mediators under inflammatory pain conditions. Future experiments will address this exciting question.

Local administration of PL2204 before induction of inflammation had a pronounced anti-inflammatory effect. This preventive effect is compatible with the observed inhibition of BK signaling mediated by the PLCβ-modulating peptide. BK is a potent inductor of plasma extravasation and hyperalgesia, and it is involved in neurogenic inflammation^60,61^. In terms of inflammation, PL2204 had a subtle effect when administered 8 days after the induction of inflammation, suggesting less participation of PLCβ at these later stages. Although certain reduction in paw swelling was found, this finding highlights the dynamism of vascular events involved in the inflammatory process. Previous anti-inflammatory effects were described after administration of the PLC inhibitor U73122^62^. However, this compound has multiple off-target effects^25–28,30,31^. Hence, our findings underscore the specific participation of PLCβ in the initial phases of inflammation.

PL2204 anti-inflammatory effect was accompanied by decrease of mechanical and thermal hypersensitivity in the injured paw during early inflammation stages. This antinociceptive activity could be due to reduced swelling, and it is in line with the decrease in TRPV1 sensitization observed in cellular studies. Our data reveal the involvement of PLCβ signaling in this mouse model of inflammatory sensitization. Previous studies found similar results in neuropathic pain models, although the PLC inhibitor was U73122 systemically administered^6^. Given the number of processes modulated by PLCβ, it is expected that unwanted effects could occur after such systemic administration. While generalized effects could be interesting for assessing the role of PLCβ in mechanistic studies, the most plausible route for potential clinical applications would be local or topical, always in the presence of pathophysiological alterations involving PLCβ recruitment.

In conclusion, peptides patterned after the autoinhibitory XY linker are effective and selective PLCβ inhibitors with *in vivo* anti-inflammatory and antinociceptive activity. PL2003 partially inhibited PLCβ enzymatic activity and BK-induced TRPV1 sensitization, being an optimal pharmacological tool for the *in vitro* study of PLCβ-dependent processes. PL2204 also showed PLCβ inhibitory activity, and its enhanced solubility in saline enables its potential use for therapeutic application. Indeed, local administration of PL2204 prevented inflammation and hypersensitivity in a mouse model of inflammatory pain. Our peptides pave the way for understanding the role of PLCβ in cellular and pathophysiological processes through the modulation of PLCβ isoforms, considered until now undruggable targets.

## ACKNOWLEDGMENTS

Financial support of the Spanish Ministry of Science, Innovation and Universities [PID2021-126423OB-C21 to A.F.C. and A.F.M. and C22 to R.G.M.] and from Generalitat Valenciana [GVA-PROMETEO/2021/031] was granted to A.F.M. J.A.L. was a recipient of an FPI fellowship from the Spanish Ministry of Science, Innovation and Universities [PRE2019-091317]. Work in N.G. laboratory was supported by the Wellcome Trust Investigator Award 212302/Z/18/Z and BBSRC Project Grant BB/V010344/1.

## AUTHOR CONTRIBUTIONS

J.A.L. executed and analyzed *in silico* experiments, designed, performed and analyzed *in vitro* assays, conducted behavioral assays and wrote the manuscript. D.C. designed and conducted behavioral assays, analyzed the data and wrote the manuscript. I.D. supervised the experiments, conceptualized the project and revised the manuscript. G.F.B. designed and supervised *in silico* experiments, and revised the manuscript. N.G. and S.S. designed and supervised PIP_2_ imaging assays and revised the manuscript, M.A.B. and R.G.M. synthesized peptides and revised the manuscript. A.F.C. conceptualized the project, provided funding and revised the manuscript. A.F.M. supervised and designed experiments, conceptualized the project, provided funding and wrote the manuscript.

## METHODS

### Building of a PLCβ3-XY linker model

To evaluate the interaction between the autoinhibitory XY linker and the isoform PLCβ3, a ligand-receptor complex was built using the structure 3OHM deposited in the Protein Data Bank (https://www.rcsb.org/). This 3D-structure describes the binding of human PLCβ3 to the alpha subunit of the mouse Gq protein, which is required for PLCβ activation. Gα_q_ protein and non-relevant molecules or ions for the activity of the XY linker were removed. Then, the crystallized region of the XY linker (575-TDEGTASSEVNATEEM-590) was isolated from the rest of the PLCβ3 structure (chain A, residues 12-471 and 593-882) and converted into an independent ligand (chain B, residues 575-590). To enable this separation, residues 591 and 592 were removed. Finally, the complex was repaired with FoldX5 to minimize its free energy by changing residues orientation with bad torsion angles, Van der Waals clashes or high total energy^63^ (https://foldxsuite.crg.eu/). Complex was built with YASARA (version 25.1.13, YASARA Biosciences GmbH, https://www.yasara.org/). The reprocessed structure of phospholipase C-β and Gq signaling complex (PDB code: 7SQ2) showed minimal differences in the catalytic domain and the XY linker (RMSD: 0.086 Å).

### Computational design of PLCβ-modulating peptides

The design of PLCβ-modulating peptides started with the fragmentation of the XY linker into smaller overlapping peptides, moving from the N-terminus to the C-terminus with an offset of 1 amino acid. Peptide-receptor complexes containing XY linker fragments ranging from 4 to 10 amino acids were generated with YASARA by removing the remaining portion of the XY loop. The binding free energy of shortened peptides was calculated with FoldX5. These values were normalized to peptide length to compare the binding of peptides of different sizes. More negative values mean better interaction with PLCβ3.

The most promising peptides, according to binding energy or proximity to PLCβ3 active site, were subjected to a sequence space search to increase their affinity. First, shortened peptide-receptor complexes were repaired with FoldX5. Then, the peptide sequence was mutated to a poly-alanine chain to later explore the fitting of the 20 natural amino acids at each position, regardless of residues in consecutive positions. After running the virtual mutagenesis, position-specific scoring matrices were generated with the binding energy of each amino acid normalized to that of the most favorable residue at each position. In these heat maps the binding energy of each residue was represented according to a color scale, with blue colors indicating a good fit, while red ones a bad fit. All computational procedures were carried out with FoldX5 (https://foldxsuite.crg.eu/).

A selection of the most energetically favorable amino acids at each position and residues with different physicochemical properties were combined to obtain designed peptides with increased theoretical affinity. Non-relevant positions for the binding of peptides were kept as in the wild-type XY linker sequence. The binding free energy of designed peptides was estimated and normalized per residue, and peptides were ranked from highest to lowest predicted affinity for PLCβ3. Non-covalent interactions between peptides and PLCβ3 were predicted with the PLIP tool (https://plip-tool.biotec.tu-dresden.de/plip-web/plip/index), using default thresholds for each interaction type^64^. Figures were drawn with open source PyMol (The PyMol Molecular Graphics System 3.0 Schrodinger, LLC, https://www.pymol.org/).

### Sequence conservation among PLCβ isoforms

Canonical protein sequences of human PLCβ1 (Q9NQ66), PLCβ2 (Q00722), PLCβ3 (Q01970) and PLCβ4 (Q15147) were downloaded from UniProt (https://www.uniprot.org/). Sequences were aligned with ClustalOmega from the European Bioinformatics Institute (EMBL-EBI, https://www.ebi.ac.uk/). Based on peptide-receptor complexes, PLCβ3 residues within 3.5 Å of peptides were identified with PyMol. Conservation of these contact residues was analyzed among PLCβ isoforms.

### Cell cultures

Human Embryonic Kidney 293 (HEK293) cells (ECACC, 85120602) were cultured in T25 cm^2^ flasks with DMEM high glucose GlutaMAX (61965-026, Gibco), supplemented with 10% fetal bovine serum (10500-064, Gibco) and 1% penicillin-streptomycin (15140-122, Gibco), at 37°C and 5% CO_2_. Upon 80% confluence, cells were split with 0.25% Trypsin-EDTA (25200-056, Gibco) for 4 min at room temperature. Trypsin reaction was stopped by adding supplemented medium, and cells were plated at the desired density on specific coated surfaces with 0.01% poly-L-lysine (P4707, Sigma-Aldrich). For calcium imaging experiments, HEK293 cells were plated onto 12 mm coverslips at 20,000 cells. For calcium microfluorimetry assays, cells were plated on 96-well black clear bottom plates (3603, Corning) at 30,000 cell/well. Cells were used 48 h after plating.

Human Embryonic Kidney 293T (HEK293T) cells (ECACC, 12022001) were cultured in T25 cm^2^ flasks with DMEM (21969-035, Gibco), supplemented with 10% fetal bovine serum (10500-064, Gibco) and 1% penicillin-streptomycin (15140-122, Gibco), at 37°C and 5% CO_2_. Upon 80% confluence, cells were split with supplemented medium. After centrifuging the cell suspension at 300 x g for 5 min at 25°C, pellet was resuspended in the appropriate volume of supplemented medium, and cells were plated at the desired density for transfection protocols.

Immortalized human keratinocyte HaCaT cells (Cytion, 300493) were cultured in T25 cm^2^ flasks with DMEM high glucose GlutaMAX (61965-026, Gibco), supplemented with 10% fetal bovine serum (10500-064, Gibco) and 1% penicillin-streptomycin (15140-122, Gibco). Upon 70% confluence, cells were washed with 0.05% PBS-EDTA for 10 min at 37°C and then detached with 0.05% Trypsin-EDTA (25300-056, Gibco) for 5 min at 37°C. Trypsin reaction was stopped by adding supplemented medium, and cells were plated onto 12 mm coverslips at a density of 20,000 cells. HaCaT cells were used 48 h after plating.

### Drugs for *in vitro* assays

PLCβ-modulating peptides were synthesized by Biomedal S.L. (Sevilla, Spain) using a solid-phase method, with the N-terminus acetylated and the C-terminus amidated. Trifluoroacetic acid was replaced by the acetate salt and purity was always above 85%. Peptides PL2001 (Ac-NATEEM-NH_2_) and PL2002 (Ac-TASSEV-NH_2_) were dissolved in PBS-1% NH_4_OH at 100 mM stock solution. Peptides PL1621 (Ac-SSEVNA-NH_2_), PL2003 (Ac-SSMTNY-NH_2_), PL2004 (Ac-NKMEMF-NH_2_), PL2204 (Ac-KRIYSSNV-NH_2_) and the randomized PL2204 (Ac-IVSNRYSK-NH_2_) were dissolved in 100% DMSO at 100 mM stock solution. Peptide PL2005 (Ac-IYSSNV-NH_2_) was prepared at 25 mM in 100% DMSO. All peptides were further diluted from the stock up to 100 µM in extracellular solution or cell medium.

U73122 hydrate (U6756, Sigma-Aldrich) was dissolved in 100% DMSO at 2 mM stock solution, which was diluted to 2 µM in extracellular solution or cell culture medium for calcium and PIP_2_ imaging assays, respectively, or to 5 µM in cell culture medium for MEA recordings. Thapsigargin (T9033, Sigma-Aldrich) was dissolved in 100% DMSO to prepare a 1 mM stock solution and diluted to 5 µM or 1 µM in assay buffer. Atropine sulphate salt monohydrate (A0257, Sigma-Aldrich) was dissolved in distilled water at 200 mM and diluted to 100 µM in assay buffer. For calcium imaging experiments, Fluo-4 AM was dissolved in 100% DMSO at 2 µg/µL. Acetylcholine (A6625, Sigma-Aldrich) was prepared in distilled water at 50 mM and diluted to 1 µM in extracellular solution. For MEA recordings, capsaicin (M2028, Sigma-Aldrich) was dissolved in 100% DMSO at 50 mM and diluted to 500 nM in extracellular solution. Bradykinin acetate salt (B3259, Sigma-Aldrich) was prepared in distilled water at 1 mM and diluted to 1 µM in extracellular solution.

### Protein expression of PLCβ3 in HEK293 cells

PLCβ3 expression was characterized in HEK293 cells by Western Blotting following a previously described protocol^65^. Total protein was extracted from HEK293 cells passage 30 in lysis buffer (150 mM NaCl, 50 mM Tris-Base, 1 mM EDTA, 1% Triton X-100, 0.5% sodium deoxycholate, 0.1% SDS, adjusted to pH 8) containing 1:100 Halt Protease Inhibitor Cocktail EDTA-free (87785, Thermo Scientific). After 40 min of incubation with the lysis buffer, solubilized proteins were collected by centrifugation at 9400 x g for 15 min. Protein concentration was determined with Pierce BCA Protein Assay Kit (23225, Thermo Scientific). 25 µg of total protein were prepared in loading buffer (60 mM Tris-HCl, 40 mM DTT, 0.01% bromophenol blue, 2% SDS, 5% glycerol, adjusted to pH 6.8) and heated up to 95°C for 5 min before loading the gel. Proteins were separated by SDS-PAGE in 7.5% polyacrylamide gels. Electrophoresis ran in running buffer (0.3% Tris-Base, 1.44% glycine, 0.1% SDS, adjusted to pH 8.4) at 100 V during the first 10 min, and then at 150 V until the end. Separated proteins were transferred onto a 0.45 µm nitrocellulose membrane (Bio-Rad) using a wet blotting system for 2 h at 100 V and 4°C in transfer buffer (25 mM Tris-Base, 190 mM glycine, 0.1% SDS, 20% methanol, adjusted to pH 8.4). Thereafter, the membrane was blocked with 5% BSA-TBST for 1 h. Isoform PLCβ3 and β-tubulin were probed with specific primary antibodies anti-PLCβ3 (1:100 in 1% BSA-TBST; sc-133231, Santa Cruz Biotechnology) and anti-β-tubulin (1:1000 in 2.5% BSA-TBST; 10094-1-AP, Proteintech) and were incubated overnight (16 h) at 4°C. After six 5-min washes with TBST (20 mM Tris-Base, 150 mM NaCl, 0.1% Tween-20, adjusted to pH 7.5), the membrane was incubated with secondary antibodies anti-mouse IgG-HRP (1:20,000 in 1% BSA-TBST; A4416, Sigma-Aldrich) to detect PLCβ3 or anti-rabbit IgG-HRP (1:25,000 in 2.5% BSA-TBST; A0545, Sigma-Aldrich) to detect β-tubulin for 1 h at room temperature. After six 5-min washes with TBST, proteins were detected with the substrate SuperSignal West Pico Plus (34577, Thermo Scientific). Chemiluminescence was visualized with the ChemiDoc imaging system (Bio-Rad).

### Calcium microfluorimetry assays

Calcium responses to acetylcholine (ACh) in HEK293 cells were characterized in a microfluorimetry assay with the Fluo-4 NW calcium assay kit (F36206, Invitrogen). 48 h after cell plating on 96-well black plates, growth medium was replaced by 90 µL Fluo-4 NW dye mix prepared in assay buffer (HBSS, 20 mM HEPES), containing 2.5 mM probenecid to prevent dye exclusion. Cells were treated with 100 µM atropine or 5 µM thapsigargin and co-incubated with the dye for 1 h at 37°C and 5% CO_2_. Cells were stabilized for 5 min at 37°C inside of the microplate reader CLARIOstar Plus (BMG Labtech GmbH, Ortenberg, Germany). Calcium transients were measured along 20 cycles spaced 150 s. After cycle 10, 100 µM ACh was automatically injected, and responses in the presence of each tested compound were measured in triplicate. Intracellular calcium responses to ACh were validated using a calcium-free assay buffer (20 mM HEPES, HBSS (14175-095, Gibco)). The fluorescent dye was excited at 485 nm and emission collected at 520 nm.

### Fluorescence calcium imaging

Non-ratiometric calcium imaging experiments were conducted with the fluorescent indicator Fluo-4 AM (F14201, Thermo Fisher Scientific). HEK293 cells were loaded with 6 µg/mL Fluo-4 AM prepared in extracellular solution (in mM: 140 NaCl, 20 D-mannitol, 10 HEPES, 5 glucose, 4 KCl, 2 MgCl_2_ and 1.8 CaCl_2_ adjusted to pH 7.4 with NaOH) containing 0.05% (w/v) Pluronic F-127 (P6867, Thermo Fisher Scientific) for 1 h at 37°C. PLCβ peptides (100 µM), vehicle or U73122 (2 µM) were co-incubated with the calcium dye for 1 h. Afterward, cells were washed with extracellular solution for at least 20 min at 37°C before starting the imaging. PLCβ was activated by applying two pulses of 1 µM ACh for 30 s, separated by a 180-s wash with extracellular solution to recover basal fluorescence levels. At the end, 1 µM ionomycin (I3909, Sigma-Aldrich) was applied to check cell integrity. Calcium imaging was conducted using an inverted fluorescence microscope (ZEIS Axiovert 200b) coupled to a Hamamatsu FLASH 4.0 LT camera (C11440-42U30, Hamamatsu, Sunayama-cho Japan). The calcium-sensitive dye was excited at 483 nm for 200 ms using the optical beam combiner Lambda 721 (Sutter Instrument, Novato, CA, USA), and fluorescence emission was collected at 512 nm every 3 s. Fluorescence intensity from individual cells was processed with the HC image DIA software (Hamamatsu Photonics).

Reponses to each stimulus were analyzed by measuring the fluorescence peak and subtracting the average basal fluorescence during the 30 s preceding stimulation. Only signals ≥ to 20 arbitrary fluorescence units were analyzed. Individual cell response to ACh was normalized to ionomycin signal. Relative ACh response was calculated by normalizing the ΔF_ACh_ / ΔF_Ionomycin_ ratio to the corresponding ratio obtained from vehicle-treated cells. The percentage of ACh-responsive cells was determined as the ratio of cells responding to ACh and ionomycin to the total number of ionomycin-responsive cells.

To assess the selectivity of PLCβ-modulating peptides, HaCaT cells were loaded with 6 µg/mL Fluo-4 AM in 0.05% (w/v) Pluronic F-127, prepared in extracellular solution (20 mM HEPES, 120 mM NaCl, 5 mM KCl, 1 mM MgCl_2_, 1 mg/mL sodium pyruvate and 1 mg/mL glucose) containing 2 mM CaCl_2_. PL2003 or PL2204 (100 µM), vehicle (0.1% DMSO) or U73122 (2 µM) were co-incubated with the Ca^2+^ dye for 1 h. Before Ca^2+^ measurements, cells were washed with 2 mM Ca^2+^-containing extracellular solution for at least 20 min. Calcium imaging recordings were initiated in Ca^2+^-free extracellular solution. Intracellular Ca^2+^ stores were depleted by 1 µM thapsigargin (TG) in 0.1 mM EGTA for 3 min. Once fluorescence signal returned to baseline, the Ca^2+^-free extracellular solution was replaced by 2 mM Ca^2+^-containing extracellular solution to induce store-operated calcium entry (SOCE). Cells responding to TG and SOCE with signals ≥ 20 arbitrary fluorescence units were included in the analysis. SOC responses were normalized to the corresponding TG signals. Relative SOC activity was calculated by normalizing the ΔF_SOC_ / ΔF_TG_ ratio to the ratio obtained from vehicle-treated cells.

### PLCδ-PH-GFP transfections and PIP_2_ imaging

HEK293T cells were plated on 24-well plates at different densities. After 24 h, those wells with approximately 60% confluence were co-transfected with a mix of 400 ng of PIP_2_ reporter PLCδ-PH-GFP encoding plasmid (Addgene plasmid #21179, gift from Tobias Meyer)^42^ and 400 ng of human bradykinin receptor B_2_R plasmid (cDNA Resource Center #BDKB200000, Rolla, MO, USA), using the transfection reagent FuGENE HD (E2311, Promega). 24 h post-transfections, cells were split and plated onto 10 mm coverslips coated with 20 µg/mL poly-D-lysine (P6407, Sigma-Aldrich). Transfected cells were cultured for 24 h before imaging.

PL2003 (100 µM), PL2204 (100 µM, 10 µM, 1 µM), vehicle or the unspecific PLC inhibitor U73122 (2 µM) were pre-incubated for 1 h before starting recordings. Basal fluorescence levels of cells were monitored for the first 30 s in an extracellular solution with calcium (in mM: 160 NaCl, 10 HEPES, 10 glucose, 2.5 KCl, 2 CaCl_2_ and 1 MgCl_2_ adjusted to pH 7.4 with NaOH). Translocation of PLCδ-PH-GFP from the plasma membrane to the cytosol was induced by a single pulse of 250 nM bradykinin (05-23-0500, Calbiochem) for 2 min. Live-cell imaging was carried out with a Nikon Eclipse TE2000-E inverted fluorescence microscope (40x objective) coupled to a IXON ultra 897 EMCCD camera (Oxford Instruments Andor, Belfast, UK). GFP was excited at 470 nm using the pE-800^fura^ Illumination System (CoolLED Ltd, Andover, UK), and fluorescence emission was collected at 535 nm every 2 s. Image analysis was done with the NIS-Elements software (Nikon Instruments Inc, NY, USA).

PLCδ-PH-GFP translocation was quantified by measuring the fluorescence increase induced by bradykinin, normalized to basal fluorescence levels (ΔF/F_0_). The time to reach maximum fluorescence intensity was evaluated from the onset of PLCδ-PH-GFP translocation. To compare the kinetics of PIP_2_ hydrolysis, fluorescence traces were normalized to their maximum and minimum values. The dose-response curve of peptide PL2204 was fitted to the following non-linear regression equation: y = a + (b - a)/(1 + (IC50/x)^hill^ ^slope^), where *y* is the % of inhibition; *x* is the peptide concentration; *a* and *b* are the minimum and maximum response, respectively; IC50 is the peptide concentration that elicits a halfway response between the maximum and the minimum, and the *hill slope* describes the steepness of the curve.

### Animals

Procedures were conducted in accordance with approval from the UMH Ethical Committee and the regional government (code: 2022 VSC PEA 0078-2), adhering to European Community guidelines (2010/63/EU) and the ethical standards of the International Association for the Study of Pain^66^. For behavioral experiments, the Institutional Animal and Ethical Committee at Universidad Miguel Hernández de Elche (UMH, Elche, Spain) approved the use of a cohort of 18 C57BL/6JRccHSd male mice (14–16 weeks old, 28–36 g; Harlan, The Netherlands), bred at the animal facility (Servicio de Experimentación Animal, UMH, Elche, Spain). For the primary culture of dorsal root ganglion neurons, neonatal Wistar rats (3-5 days-old, Harlan, The Netherlands) were obtained from the breeding stock at UMH. All animals were housed under controlled conditions (22 ± 1°C, 55 ± 20% relative humidity) with a 12-hour light/dark cycle (lights on from 8:00 a.m. to 8:00 p.m.). Efforts were made to habituate the animals to handling and to minimize pain and stress.

### Primary culture of DRG neurons

Dorsal root ganglion neurons (DRGs) were isolated from neonatal Wistar rats (3-5 days-old), humanely euthanized by decapitation. DRGs were harvested from all the spinal levels, with nerve fibers removed, and enzymatically digested for 1 h at 37°C and 5% CO_2_ with a 2.5 mg/mL collagenase type IA solution (C9891, Sigma-Aldrich) in DMEM high glucose GlutaMAX (10566-016, Gibco) medium supplemented with 1% penicillin-streptomycin (15140-122, Gibco). Thereafter, ganglia were mechanically dissociated using a 1 mL micropipette. Single cell suspension was filtered through a 100 µm cell strainer and sensory neurons washed after three successive centrifugations at 300 x g for 5 min in DMEM high glucose GlutaMAX medium supplemented with 1% penicillin-streptomycin and 10% fetal bovine serum (10500-064, Gibco). Finally, DRG neurons were plated in drops on multielectrode array chambers coated with 8.3 µg/mL poly-L-lysine (P9155, Sigma-Aldrich) and 5 µg/mL Cultrex 3-D Culture Matrix Laminin I (3446-005-01, Bio-Techne R&D Systems). After 30 min allowing cell adhesion, cultures were supplemented with 50 ng/mL 2.5S nerve growth factor (N6009, Sigma-Aldrich) and 0.6 µg/mL cytosine arabinoside (C1768, Sigma-Aldrich) in the complete medium. Neuronal cultures were maintained at 37°C and 5% CO_2_ for 48 h before starting recordings.

### Multielectrode array recordings

Extracellular action potentials were recorded in thin multielectrode planar arrays containing 60 electrodes (59 recording electrodes plus one internal reference electrode) with a diameter of 30 µm and an interelectrode distance of 200 µm. The electrical activity was recorded with the MEA1060-INV System at a sampling frequency of 25 kHz, controlled by MC_Rack software version 4.6.2 (Multi Channel Systems MCS GmbH, Reutlingen, Germany). Recordings were performed at 34.5°C with the temperature controller TCO2 (Multi Channel Systems MCS GmbH). PLCβ peptides (100 µM) or vehicle (0.1% DMSO) were pre-incubated for 1 h in DMEM high glucose GlutaMAX medium supplemented with 10% fetal bovine serum and 1% penicillin-streptomycin, in absence of nerve growth factor. The effect of the unspecific PLC inhibitor U73122 was also tested at 5 µM. To evaluate TRPV1 sensitization, three pulses of 500 nM capsaicin (P1, P2 and P3) were applied for 30 s. Between P1 and P2, cells were washed for 150 s with extracellular solution (in mM: 140 NaCl, 20 D-mannitol, 10 HEPES, 5 glucose, 4 KCl, 2 MgCl_2_ and 1.8 CaCl_2_, adjusted to pH 7.4 with NaOH). Between P2 and P3, after a 120-s wash, the neuronal culture was stimulated with 1 µM bradykinin for 8 min to activate the PLCβ pathway and potentiate TRPV1. At the end, neuronal excitability was assessed by applying a short pulse of 40 mM KCl. Stimuli were applied using a peristaltic perfusion system PPS2 (Multi Channel Systems MCS GmbH) at a flow rate of 5 mL/min.

Recordings were analyzed using MC_Rack software (Multi Channel Systems MCS GmbH). Raw data were filtered with a second order high pass Butterworth filter, using a cutoff frequency of 50 Hz to remove background noise. Spikes were detected in the filtered data when their amplitude exceeded a threshold automatically set for each electrode at ± 5 µV the standard deviation of the recording. Mean spike frequency for each stimulus was calculated in the 60-s time interval upon stimulation. Only KCl-responsive electrodes were included in the analysis. TRPV1 sensitization was quantified by comparing responses to capsaicin before (P2) and after applying bradykinin (P3) through the relation P3 - P2. The percentage of BK response or TRPV1 sensitization under each treatment was determined by normalizing the corresponding values to those of vehicle.

### Experimental design of behavioral experiment

To maximize the information obtained from the authorized number of animals, the cohort of 18 mice was divided into three groups of six (**Figure S5**). Two groups were exposed to inflammation (i.pl. CFA, n=6 each), while the third served as a control (i.pl. saline, n=6).

To evaluate the preventive effect of PL2204 on inflammation and hypernociception, one of the CFA-exposed groups received vehicle (5% DMSO in saline), and the other was pre-treated with PL2204. Researchers were blinded to the treatment in the CFA-treated groups (vehicle-CFA vs. PL2204-CFA) but unblinded for the control group. To assess possible therapeutic effects of the peptide, when both CFA-treated groups displayed similar behavior (8 days post-CFA injection), the vehicle-treated group was switched to PL2204, while the PL2204-treated group received vehicle (**Figure S5**). The control group, which did not receive CFA, was injected with vehicle in the same manner.

### Model of Inflammation

Inflammation and the associated nociceptive sensitization were induced by subcutaneous injection of 10 µL of 0.5 mg/mL CFA (Complete Freund’s Adjuvant, F5881, Sigma Aldrich, USA) in saline into the plantar side of the left hind paw using a Hamilton syringe with a 30-gauge needle. Control animals received 10 µL of saline.

### Drugs tested in behavioral experiments

PL2204 was dissolved in 5% DMSO in saline at 10 mg/mL. A single dose of PL2204 (25 µL, 250 µg) was administered via a 0.3 mL syringe (324826, BD Micro-Fine Demi, BD Medical, France) into the plantar side of the left hind paw, either 2 h before (preventive effect) or 8 days after (curative effect) the induction of inflammation. Control mice received 5% DMSO in saline.

### Paw inflammation assessment

Paw thickness was measured before and at 2 h, 24 h, and 4, 7, 8, and 9 days after CFA or saline injection. A manual caliper with a precision of 0.02 mm was used, always at the same location between the insertions of the 1st and 5th digits.

### Antinociceptive evaluation

<u>Mechanical sensitivity</u> was assessed by evaluating the hind paw withdrawal thresholds on days −3, −2, −1, and at 3 h, 24 h, and 4, 8, and 9 days after CFA injection. Mice were placed in Plexiglas® chambers (10 × 10 × 14 cm) with a wire grid bottom and habituated for 1 h before testing. Von Frey filaments equivalent to 0.04, 0.07, 0.16, 0.4, 0.6, 1 and 2 g were used, applying first the 0.4 g filament and increasing or decreasing the strength according to the response^67^. Four additional filaments were sequentially applied since the first change of response (from negative to positive or from positive to negative). The sequence of the last six responses was used to calculate the withdrawal threshold following the method described by Dixon^68^.

<u>Heat Sensitivity</u> was assessed using a plantar test apparatus^69^ (Ugo Basile, Italy). Animals were habituated for 1 h before testing. Radiant heat was applied to the plantar surface of the hind paw, with the beam intensity adjusted to achieve baseline latencies of 8–12 seconds in control mice. A cutoff time of 15 seconds was applied. The mean hind paw withdrawal latencies were obtained from the average values of three separate trials, taken at 5- to 10-min intervals, to reduce the possible influence of thermal sensitization on the response.

### Statistical analyses

For cellular studies, data normality was first assessed with Shapiro-Wilk test. Depending on the distribution, statistical significance between two groups was determined with an unpaired Student’s t-test or the Mann-Whitney U test. Statistical significance between more than two groups was assessed with a one-way ANOVA followed by Bonferroni’s *post hoc* test or a Kruskal-Wallis followed by Dunn’s test. Data points were considered outliers and removed if they exceeded 1.5 times the interquartile range (Q3 – Q1) above the third quartile (Q3) or below the first quartile (Q1). The number of data points per group is shown in brackets, while the number of biological replicates (N) is indicated in the figure legends. For the *in vivo* experiment, a two-way ANOVA or a mixed model (within-factor: "time"; between-factor: "treatment", and their interaction) were used, followed by *post hoc* Tukey tests to compare groups and by Dunnett’s tests to compare vs. baseline. In the manuscript, data are expressed as mean + Standard Error of the Mean, while statistical significance was set at p<0.05. Statistical analyses were done with GraphPad Prism 9.4.1 (GraphPad Software Inc., San Diego, CA, USA). Source data and statistical analyses for each figure are provided in the supplementary file Source_Data.zip.

## SUPPLEMENTARY FIGURE LEGENDS

**Figure S1.**
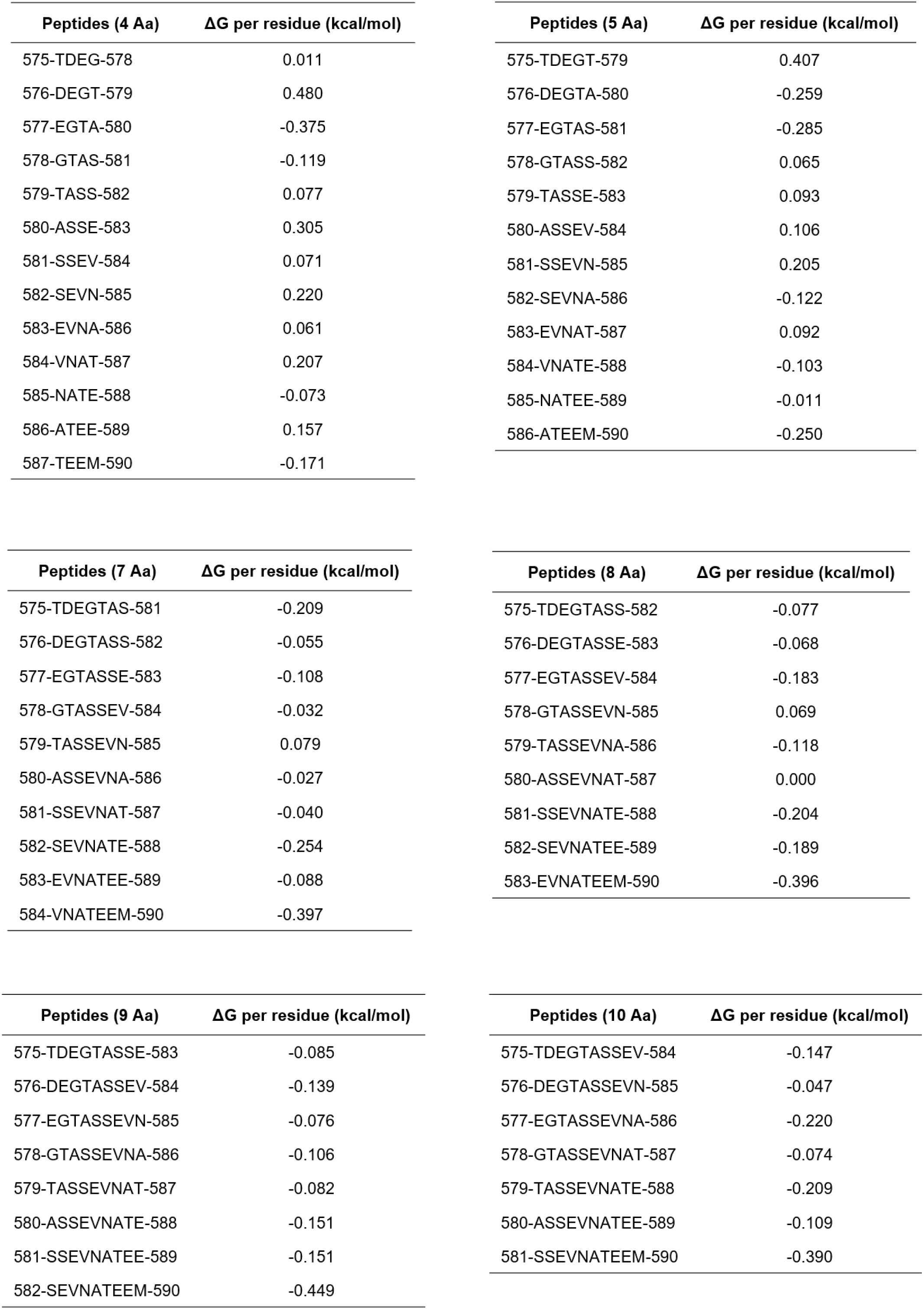
ΔG per residue of peptides containing 4, 5, 7, 8, 9 and 10 amino acids obtained after the fragmentation of the XY linker. More negative value means better interaction. ΔG: binding free energy. Aa: amino acid.

**Figure S2.**
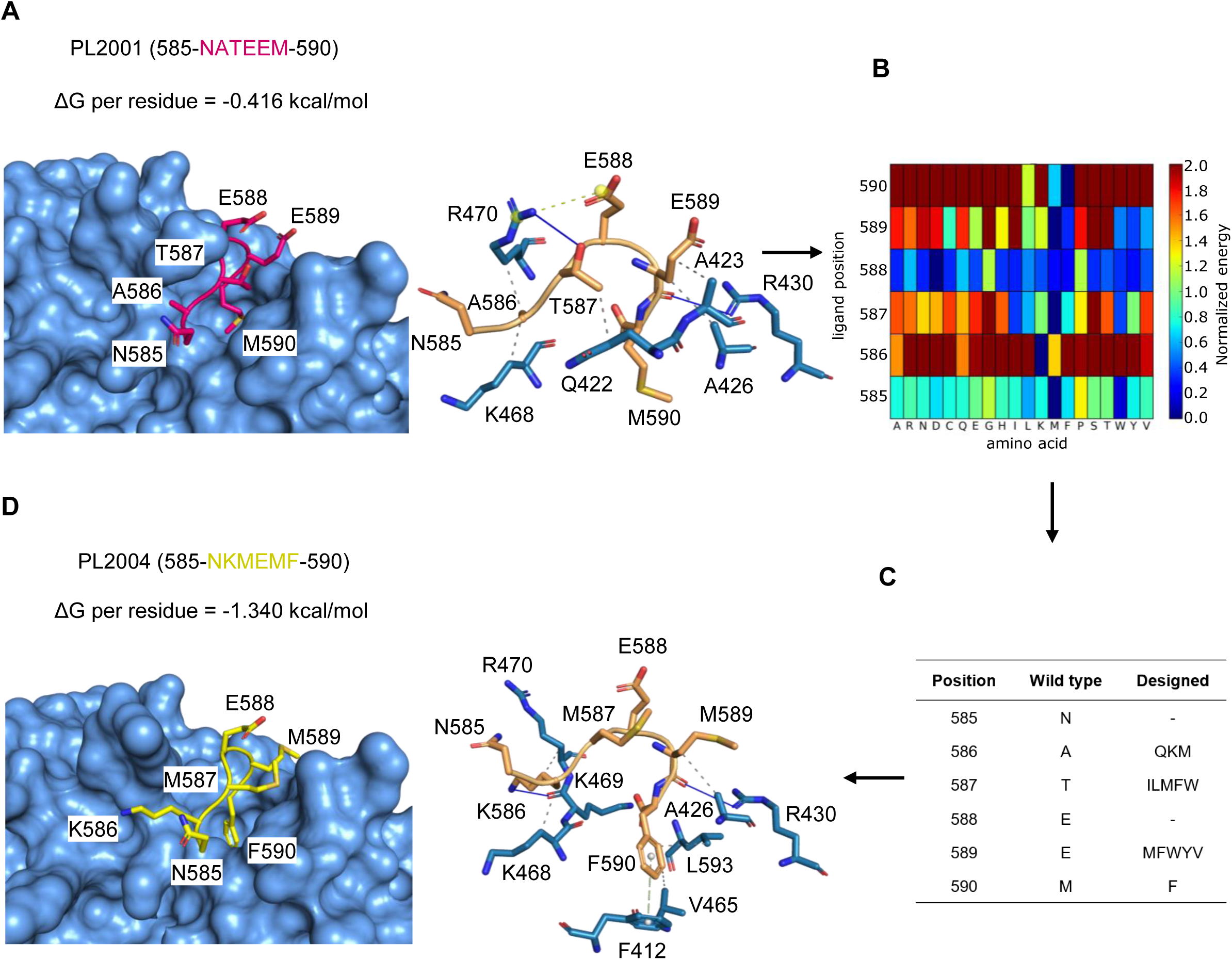
PLCβ peptide 585-NATEEM-590 patterned after the autoinhibitory XY linker with the best binding free energy. **(A) Left panel,** anchor points of wild-type peptide 585-NATEEM-590 (pink, PL2001) bound to PLCβ3 isoform with predicted ΔG per residue. **Right panel**, PLCβ3 residues directly interacting with this peptide are in blue. Solid blue lines: hydrogen bonds. Dashed yellow lines: salt bridges. Dashed grey lines: hydrophobic interactions. **(B)** Sequence space search to optimize interaction of wild-type peptide with PLCβ3. Heat maps shows normalized binding energies of the 20 natural amino acids (x-axis) at each position (y-axis) according to a color scale from blue (best fit) to red (worst) **(C)** The most favorable residues (Designed column) at each position (Position column) of the wild-type peptide (Wild type column) were combined to obtain new peptides. **(D) Left panel,** designed peptide 585-NKMEMF-590 (yellow, PL2004) shows increased theoretical affinity compared to wild-type (ΔG per residue). **Right panel**, PLCβ3 residues directly interacting with this peptide are in blue. Solid blue lines: hydrogen bonds. Dashed green lines: π-stacking. Dashed grey lines: hydrophobic interactions. ΔG: binding free energy.

**Figure S3.**
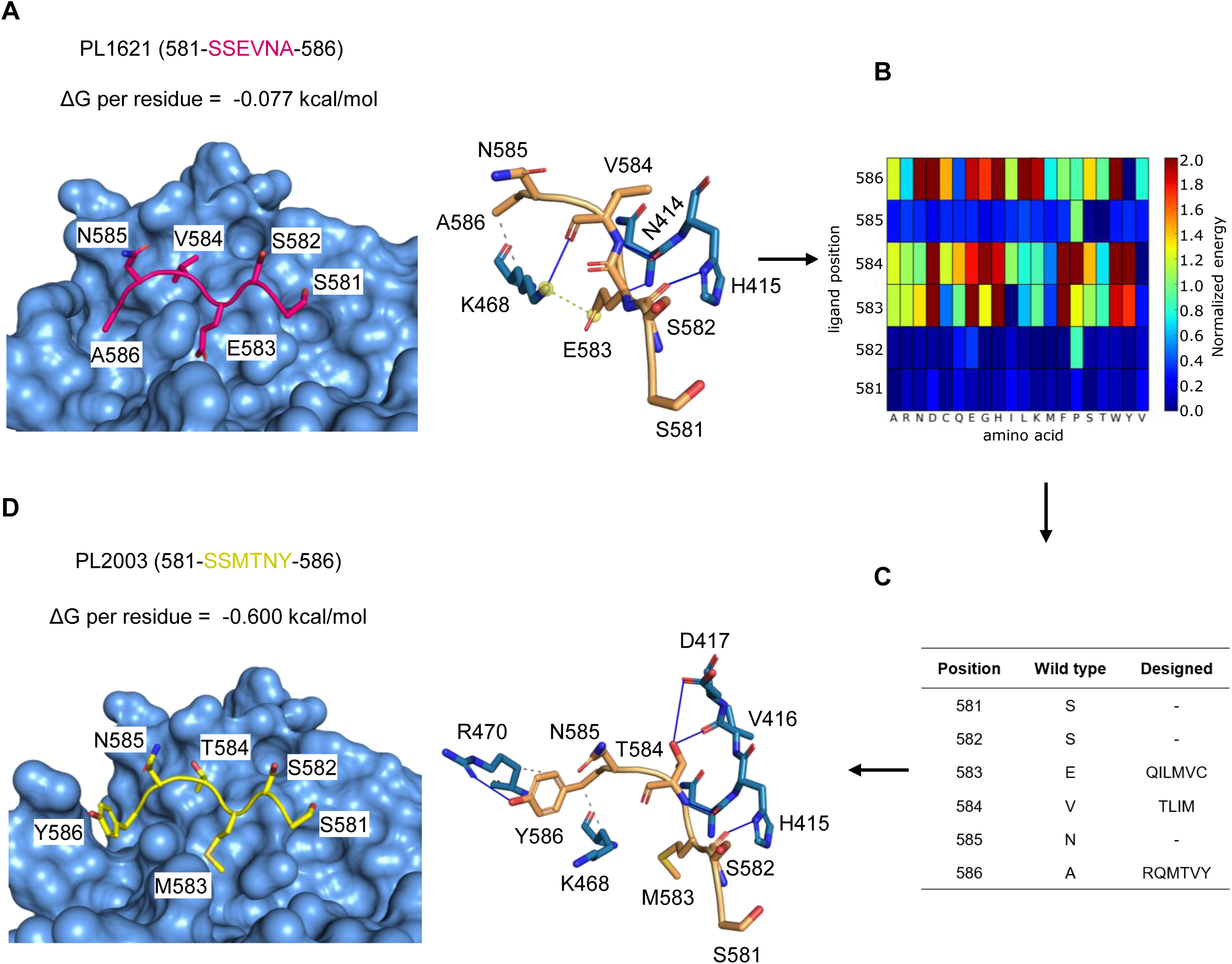
PLCβ peptide 581-SSEVNA-586 patterned after the lid helix region of the autoinhibitory XY linker. **(A) Left panel**, anchor points of wild-type peptide 581-SSEVNA-586 (pink, PL1621) bound to PLCβ3 isoform with predicted ΔG per residue. **Right panel**, PLCβ3 residues directly interacting with this peptide are in blue. Solid blue lines: hydrogen bonds. Dashed yellow lines: salt bridges. Dashed grey lines: hydrophobic interactions. **(B)** Sequence space search to optimize interaction of wild-type peptide with PLCβ3. Heat map shows normalized binding energies of the 20 natural amino acids (x-axis) at each position (y-axis) according to a colour scale from blue (best fit) to red (worst fit). **(C)** The most favorable residues (Designed column) at each position (Position column) of the wild-type peptide (Wild-type column) were combined to obtain new peptides. **(D) Left panel**, designed peptide 581-SSMTNY-586 (yellow, PL2003) shows increased theoretical affinity compared to wild-type (ΔG per residue). **Right panel**, PLCβ3 residues directly interacting with this peptide are in blue. Solid blue lines: hydrogen bonds. Dashed grey lines: hydrophobic interactions. ΔG: binding free energy.

**Figure S4.**
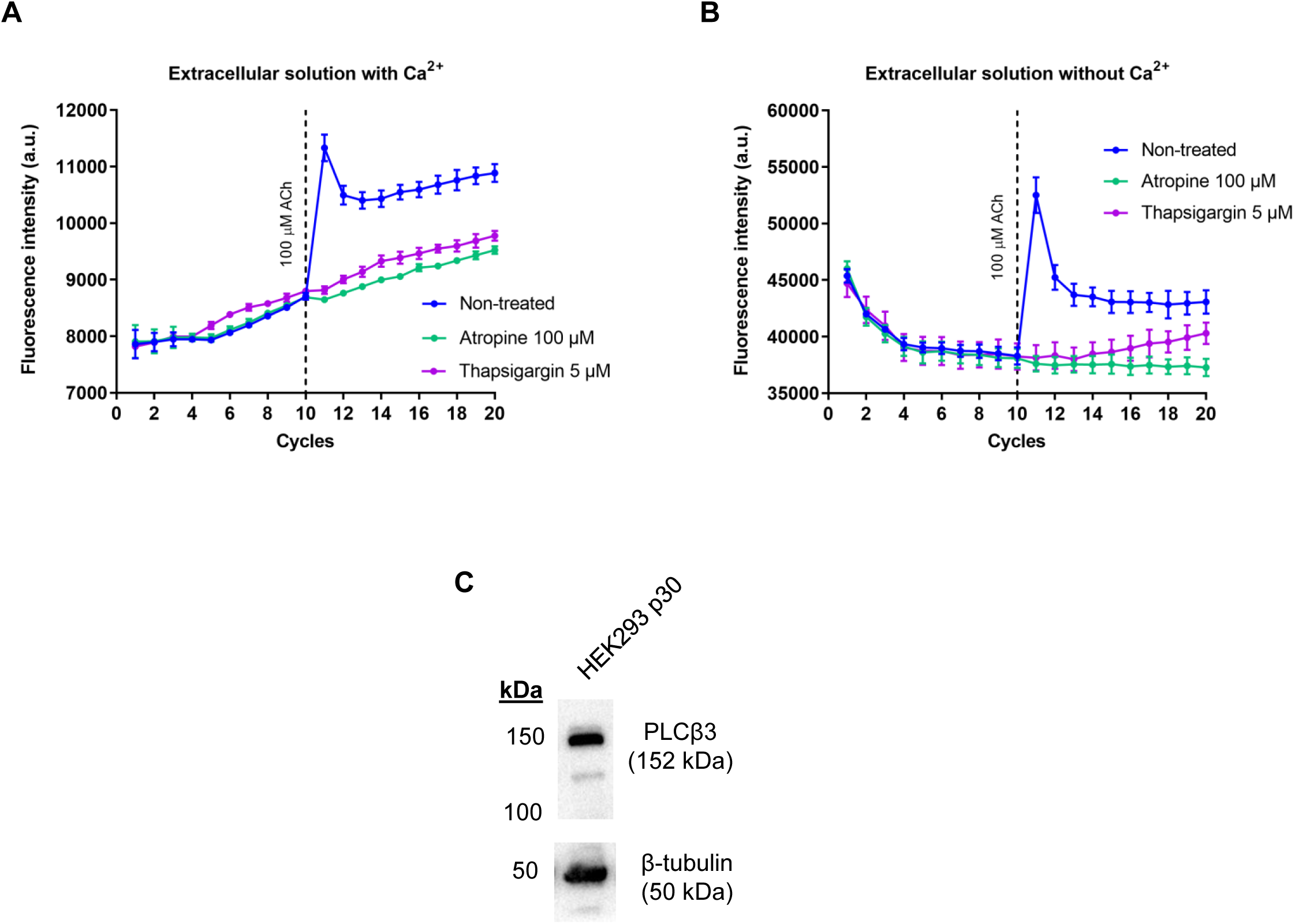
ACh in HEK293 cells activates muscarinic receptors coupled to PLCβ. **(A)** Pre-incubation of HEK293 cells with the muscarinic antagonist atropine or the Ca^2+^-ATPase inhibitor thapsigargin abolishes ACh-induced Ca^2+^ transients. Fluorescence emission was measured 20 times (cycles) in Ca^2+^ microfluorimetry assays, using an extracellular solution with Ca^2+^ with the dye Fluo-4. ACh was injected after cycle 10. **(B)** As in A, but using a Ca^2+^-free extracellular solution. **(C)** Western blot of PLCβ3 (152 kDa) and loading control β-tubulin (50 kDa) in whole protein lysate from HEK293 cells passage number 30. **(A,B)** Data points represent the mean ± Standard Error of Mean of n=3 technical replicates. ACh: acetylcholine.

**Figure S5.**
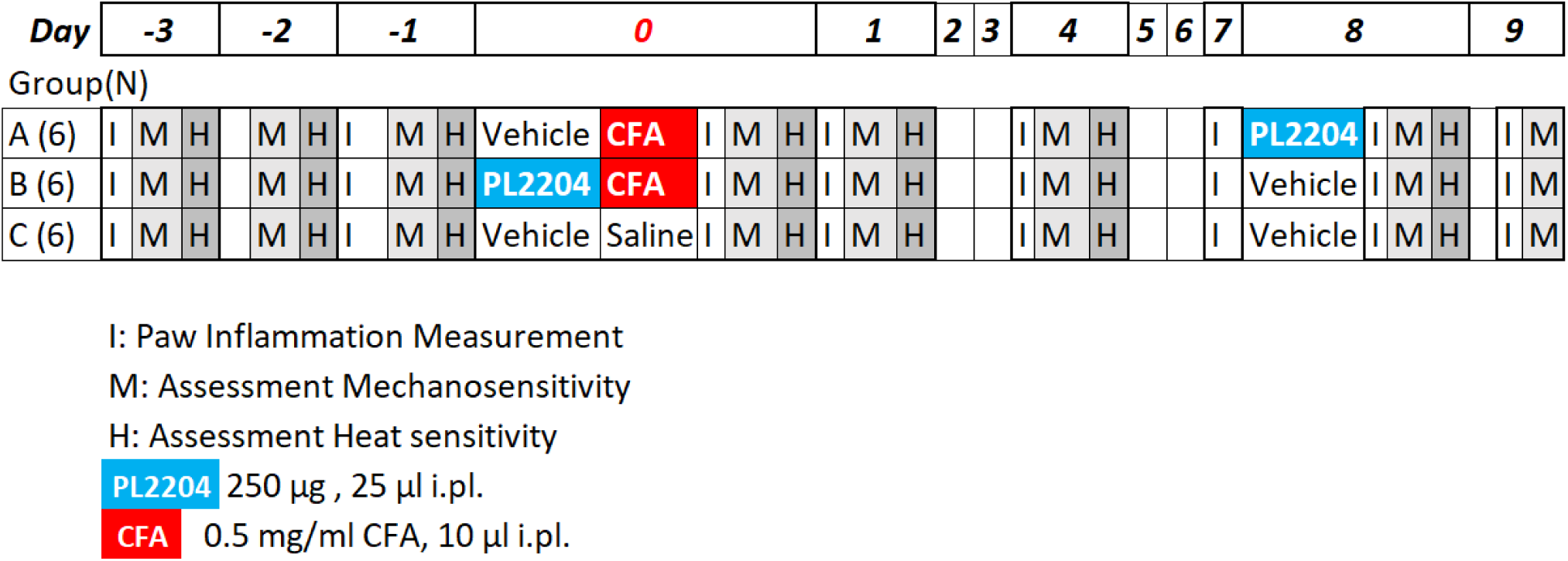
Experimental design of behavioral experiment. Mice were distributed in 3 homogeneous groups according to their mechanical sensitivity and paw thickness. Two of the groups were injected with CFA, and of those, one was pre-treated with vehicle **(A)** and the other with PL2204 **(B)**. The remaining group **(C)** received saline injection in the paw and vehicle pre-treatment. By day 8, when CFA-treated groups had similar levels of inflammation, PL2204 was administered to the CFA-treated group that was never exposed to PL2204 **(A)**, whereas the group previously exposed to PL2204 received vehicle **(B)**. The control group received vehicle at this point. Paw edema (I) was measured on days −3,-2,-1, and 2 h, 24 h and 4, 7, 8 and 9 days after CFA injection. Mechanical sensitivity (M) was measured on days −3,-2,-1, and 3 h, 24 h and 4, 8 and 9 days after CFA injection, while heat sensitivity (H) was measured on days −3,-2,-1, and 3 h, 24 h and 4 and 8 days after CFA injection. CFA: complete Freund’s adjuvant. i.pl.: intraplantar.

**Figure S6.**
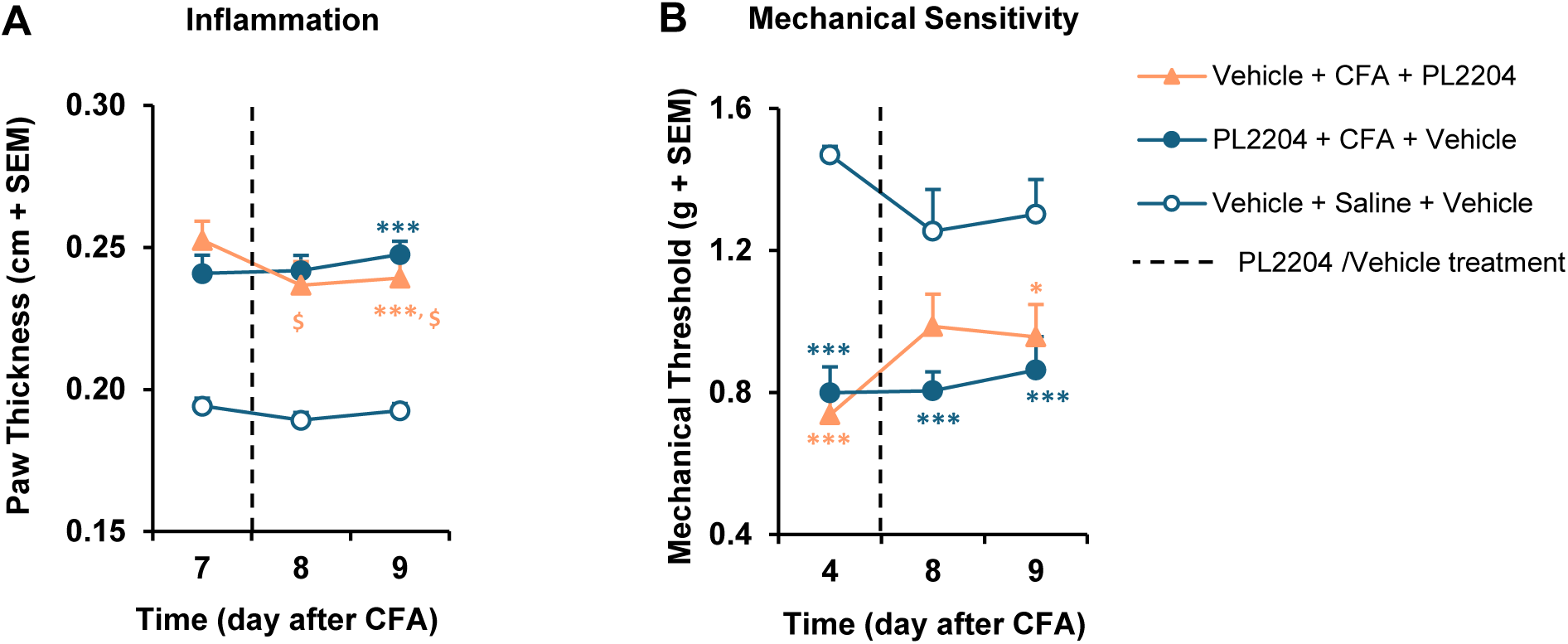
Effect of a local curative treatment with PL2204 after CFA-induced inflammation. **(A)** Mice treated with PL2204 8 days after CFA experienced a slight decrease of inflammation (vs. pre-treatment value on day 7), although swelling was still pronounced ( vs. Vehicle+Saline+Vehicle). **(B)** Mice treated with PL2204 8 days after CFA showed partial alleviation of mechanical hypersensitivity (N.S. vs. Vehicle+Saline+Vehicle; N.S. vs. mice receiving PL2204+CFA+Vehicle). **(A,B)** Mean + Standard Error of Mean (SEM) values are shown. Two-way Anova followed by Tukey and Dunnett’s tests. *p<0.05, ***p<0.001 vs. control (Vehicle+Saline+Vehicle). ^$^p<0.05 vs. baseline. N=6 animals per group. CFA: complete Freund’s adjuvant.

**Figure S7.**
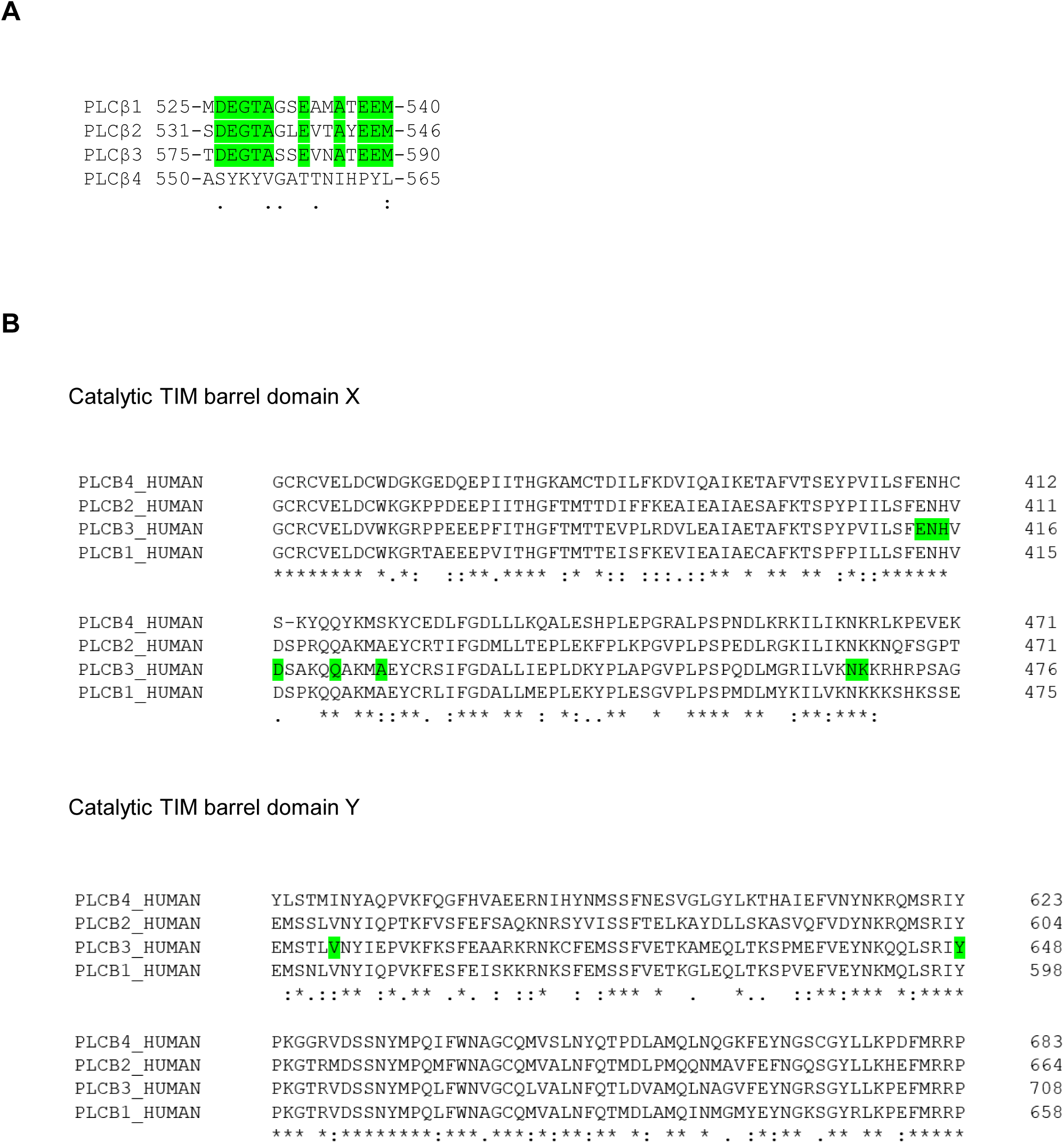
PLCβ-modulating peptides patterned after the XY linker of PLCβ3 interact with a highly conserved region among PLCβ isoforms. **(A)** Multiple sequence alignment of PLCβ isoforms focused on the crystallized region of the XY linker in the PLCβ3 structure. PLCβ4 shows a higher sequence divergence with the rest of isoforms. Conserved residues among PLCβ1, β2 and β3 isoforms are highlighted in green. ( : ) Conservation between groups of strongly similar properties. ( . ) Conservation between groups of weekly similar properties. ( ) Not conserved. **(B)** Multiple sequence alignment of PLCβ isoforms focused on regions of the catalytic TIM barrel domain that interact with PLCβ-modulating peptides. Positions highlighted in green indicate PLCβ3 residues within 3.5 Å of peptides. These residues are mostly conserved among PLCβ isoforms. ( * ) Fully conserved residue. ( : ) Conservation between groups of strongly similar properties. ( . ) Conservation between groups of weekly similar properties. ( ) Not conserved.

